# Whole genome duplication uncouples cell volume scaling from mitochondrial homeostasis

**DOI:** 10.64898/2026.10.28.762444

**Authors:** Anna Regina Hertel, Araba Mansa Sagoe, Galal Yahya, Anna-Lena Heusser, Azkia Khan, Markus Räschle, Johannes M. Herrmann, Zuzana Storchová, Prince Saforo Amponsah

**Author notes:** Corresponding authors: Prince Saforo Amponsah, Zuzana Storchova. These authors contributed equally to the work (Anna Regina Hertel, Araba Mansa Sagoe).

## Abstract

Whole genome duplication, or polyploidy, increases cell size and nuclear volume, yet how cells maintain cytonuclear balance and metabolic fitness following ploidy expansion remains unclear. Using isogenic series of *Saccharomyces cerevisiae* strains ranging from haploid (1N) to tetraploid (4N), and isogenic human diploid and tetraploid cell lines, we investigated the impact of polyploidy on mitochondrial homeostasis and metabolic adaptation. Quantitative proteomics uncovered a sub-linear scaling of mitochondrial proteins with increasing ploidy, with pronounced depletion of respiratory chain components, mitochondrial DNA maintenance factors, and redox homeostasis proteins. These changes were highly conserved, indicating a conserved consequence of genome expansion. Impaired mitochondrial scaling reduced respiratory growth, membrane potential and oxygen consumption, and elevated oxidative stress. In budding yeast, the mitochondrial dysfunction triggered reactive oxygen species (ROS) accumulation and an unconventional activation of stress-responsive signaling pathways, including the Hog1 and cell integrity MAPK networks, which converged on Rtg2-dependent retrograde signaling to promote mitochondrial metabolism despite chronic dysfunction. Our findings identify defective mitochondrial scaling as a key consequence of polyploidy and establish ROS- and stress signaling-dependent retrograde response as a key adaptive mechanism linking genome duplication to mitochondrial homeostasis and cellular fitness.

## Introduction

Most eukaryotic organisms maintain a diploid (2N) life-cycle, meaning that they contain two sets of nearly identical, homologous chromosomes. An evolutionary advantage of diploidy is in masking deleterious mutations, while effective purifying selection is retained during meiosis with haploid gametes^1^. While diploidy is most common, changes in ploidy are observed frequently in nature. Increased ploidy of the entire organism through fertilization with unreduced gametes, or whole genome duplication (WGD) through changes in cell division cycle increases the number of available gene copies. This provides genetic redundancy, enabling mutations to accumulate in duplicate genes without detrimental effects. Moreover, WGD can drive evolution as it promotes the development of novel gene functions as well as the separation of existing functions^2^. Thus, increased ploidy is often associated with increased stress resistance and elevated adaptability.

Frequently, however, genome doubling is a detrimental event. Acute WGD profoundly perturbs cellular physiology, affecting cell cycle progression, chromosome segregation, genome stability, metabolism, and stress responses^3–7^. The sudden increase in genome content is accompanied by extensive cellular enlargement, imposing scaling constraints on the proteome and intracellular organelles, which challenge the maintenance of homeostasis. Consequently, nascent polyploids commonly experience reduced fitness, proteome imbalance, oxidative and proteotoxic stress, and impaired proliferation, creating strong selective pressure for adaptive mechanisms that restore cellular and metabolic equilibrium^8^.

One of the most prominent consequences of polyploidy is an increase in cell size, likely due to the expansion of genome content and the accompanying rise in biosynthetic and energetic demands. In budding yeast, cell volume increases nearly proportionally with genome copy number, approximately doubling after each whole genome duplication event^9^. It should be noted, however, that cell volume increases more rapidly than surface area. Thus, polyploidization reduces the surface-to-volume ratio, thereby altering the physical and metabolic constraints governing cellular physiology. Consistent with this, polyploid cells exhibit impaired nutrient uptake, reduced waste exchange, and slower proliferation, while enlarged cells frequently experience elevated oxidative stress and genome instability^10^. Similar scaling effects have been reported in polyploid amphibians, where increased ploidy is associated with reduced oxygen consumption, potentially reflecting lower energetic costs for maintaining plasma membrane potential due to the decreased membrane area relative to cytoplasmic volume^8^. Although cell size and total mRNA abundance generally scale proportionally with genome content, protein abundance scales sub-linearly – meaning that protein abundance increases less than expected relative to cell volume –, indicating that proteome expansion does not fully match cellular growth^11, 12^. Importantly, increasing cell size alters the morphology, abundance, and physiology of intracellular organelles in both budding yeast and mammalian cells^12, 13^. While the consequences of genome duplication for chromosome segregation and genome stability have been extensively investigated (e.g., ^14^), much less is known about how mitochondrial content and function scale with polyploid cell growth. Given the central role of mitochondria in energy production, redox homeostasis, and biosynthetic metabolism, understanding how mitochondrial homeostasis is coordinated with increased cell size is likely to reveal fundamental constraints governing the physiology and fitness of polyploid cells.

How whole genome duplication influences the behavior and function of cellular organelles, especially mitochondria, in eukaryotic cells, is an intriguing and understudied question. Although the vast majority of mitochondrial proteins (about 900 in yeast and 1,300 in humans) are encoded by nuclear genes and subsequently imported, 8 and 13 proteins in yeast and humans, respectively, remain encoded by the mitochondrial genome. Because these proteins are of crucial relevance in respiring organisms and assemble into multisubunit complexes, any imbalance between nuclear- and mitochondrially-encoded components can disrupt complex formation, compromise respiratory efficiency, and trigger proteostatic and energetic stress^15–17^. Moreover, duplicating the entire nuclear genome inherently elevates cellular energy requirements, placing additional demand on mitochondria biogenesis, import machinery, and bioenergetic capacity. With increasing ploidy, the mitochondrial DNA (mt-DNA) copy number roughly scales up with ploidy, with diploid strains having approximately twice the number of mitochondrial genomes and triploid having nearly three times the number, however, tetraploid and pentaploid cells show a clear sub-scaling of mitochondrial genomes^18^. Accordingly, an increasing ploidy can cause detrimental effects on mitochondria even though the mechanistic details for the genetic interactions of the two genomes remain largely unclear.

Mitochondrial network size and mt-DNA are dynamically regulated during cell proliferation, with mitochondrial mass typically increasing approximately twofold during each cell cycle to support the energetic and biosynthetic demands of daughter cell growth^19–22^. Therefore, genome duplication and the associated increase in cell size would be expected to require a corresponding expansion of mitochondrial content. Mitochondrial abundance is determined by the balance between mitochondria biogenesis and selective removal of damaged organelles through mitophagy, while mitochondrial morphology and functional capacity are shaped by the opposing processes of fission and fusion^23–25^. However, whether mitochondrial scaling remains proportional following changes in genome content remains poorly understood. Quantitative analyses in proliferating diploid mammalian cells have revealed nonlinear relationships between mitochondrial function and cell size, as measured by membrane potential-dependent dye uptake, with maximal mitochondrial activity observed at intermediate cell sizes and declining function in larger cells^26, 27^. In contrast, studies in budding yeast have suggested that mitochondrial network size scales approximately linearly with cell growth, indicating that mitochondrial scaling may be tightly coordinated under normal physiological conditions^21^. At the molecular level, mitochondria biogenesis is integrated with cellular growth through nutrient-sensing pathways, particularly the mTOR signaling network, which couples anabolic activity, biomass accumulation, and organelle expansion^28–30^. Whether these regulatory mechanisms are sufficient to maintain mitochondrial homeostasis following whole genome duplication remains an important unanswered question.

From a simplistic viewpoint, mitochondrial metabolism is primarily driven by oxidative phosphorylation (OXPHOS), which generates energy in the form of ATP via the respiratory chain, and the tricarboxylic acid (TCA) cycle, which supplies metabolic intermediates to feed OXPHOS, amino acid, lipid and nucleotide biosynthesis^31^. A tightly regulated crosstalk between the nucleus and mitochondria on the metabolic or energy state of the cell directs mitochondrial metabolism. Under normal physiological conditions the nucleus mediates transcription of genes that drive the TCA cycle and OXPHOS in response to low ATP and amino acid levels, in a process termed anterograde signaling^32^. However, compromised mitochondrial health (e.g., persistent drop in membrane potential^33^ or mt-DNA depletion^34^) or impaired respiratory functions elicits the reverse communication from mitochondria to nucleus, which enables the transcription of genes to bypass respiratory deficiency and ensure biosynthetic processes occur via the TCA cycle to sustain cell growth and proliferation, in a mechanism called retrograde signaling^35, 36^. Thus, the retrograde pathway rewires cellular metabolism in adaptation to chronic mitochondrial dysfunctions by reprogramming nuclear gene expression^31^. Mitochondrial retrograde signaling occurs in all eukaryotes, but the molecular players are diverse^35^. This mechanism is distinguishable from responses to acute mitochondrial stress, like the unfolded protein (UPR^mt^) and integrated stress (ISR) responses, which are traditionally triggered by mitochondrial protein import and folding problems and reactive oxygen species (ROS)^37–39^. In eukaryotes, both anterograde and retrograde signaling are under the control of protein kinases and phosphatases including AMPK, MAPK, and mTOR^40–43^. In budding yeast, canonical retrograde signaling is a transcriptional response governed by the RTG system; Rtg2 is the main cytosolic effector, while Rtg1/3 are transcriptional factors that shuttle between the cytoplasm and nucleus. If, and how, retrograde signaling contributes to the resilience and proliferation of eukaryotic cells with chronic whole genome duplication, is a largely underexplored topic.

Here, we set out to evaluate how increasing ploidy affects mitochondria in eukaryotic cells. By analyzing series of isogenic yeast strains and human cell lines of different ploidy, we show that the mitochondrial proteome subscales with ploidy and that mitochondrial function becomes compromised as a result. We find that Rtg2, a pivotal mitochondrial retrograde signaling factor in *Saccharomyces cerevisiae*, becomes stabilized in response to increased ploidy, and the abundance of its downstream targets increase, constituting a cellular distress signal that reflects underlying mitochondrial dysfunction. Our work outlines mechanisms by which genome duplication perturbs mitochondrial homeostasis and engages the retrograde response to rescue organellar imbalance.

## Results

### Polyploidy remodels the mitochondrial sub-proteome and impairs yeast growth under respiratory conditions

The size or volume of eukaryotic cells scales linearly with their genomic content or ploidy^9, 11^. Changes in cell size are accompanied by a proportional change in the global proteome, proteomic content of organelles, organellar size and distribution^12, 13, 21, 22, 44^. Analysis of the proteomic content of our previously generated isogenic series of budding yeast with different ploidy, from haploid to tetraploid, revealed that while cell size scales linearly with ploidy, the global proteome scales sub-linearly^11^. To understand how the scaling impacts the proteome content of cellular organelles, we analyzed the size scaling characteristics of each measured protein by calculating their slope values, which describe how the relative concentration of each protein changes with cell size^12^. We then sorted the proteins according to their subcellular localization. This revealed that various proteins show different scaling behavior (Fig. 1a). While the mean slope of proteins localized to the nucleolus, nucleus and cytoplasm scaled similarly or slightly higher than the global proteome, the mitochondrial proteins adopted a sub-scaling behavior (mean slope; global proteome: 0.08, nucleolus: 0.09, nucleus: 0.13, cytoplasm: 0.18, mitochondrial: -0.1) (Fig. 1a), suggesting that polyploidy-induced cell size increase strongly dilutes the mitochondrial sub-proteome. Indeed, the median fold change in abundance of mitochondrial proteins (M) diminishes with increasing ploidy even in comparison with the global proteome (P) (median fold change; **2N**: P = 1.61, MP = 1.51; **3N**: P = 2.31, MP = 2.19; **4N**: P = 2.95, MP = 2.83) (Fig. 1b). Although mitochondrial proteins sub-scale on average with increasing ploidy, there is a striking difference among individual proteins and specific functional groups, as revealed by analysis of the log2-fold changes across ploidy series. Particularly, the abundance of proteins belonging to the respiratory/electron transport chain (ETC), mt-DNA maintenance and the antioxidant response groups was consistently and significantly reduced, while mitochondrial ribosomal subunits and import machinery components were variably deregulated, and TCA cycle proteins were significantly upregulated in polyploid yeasts (Supplementary Fig. 1a).

**Figure 1:**
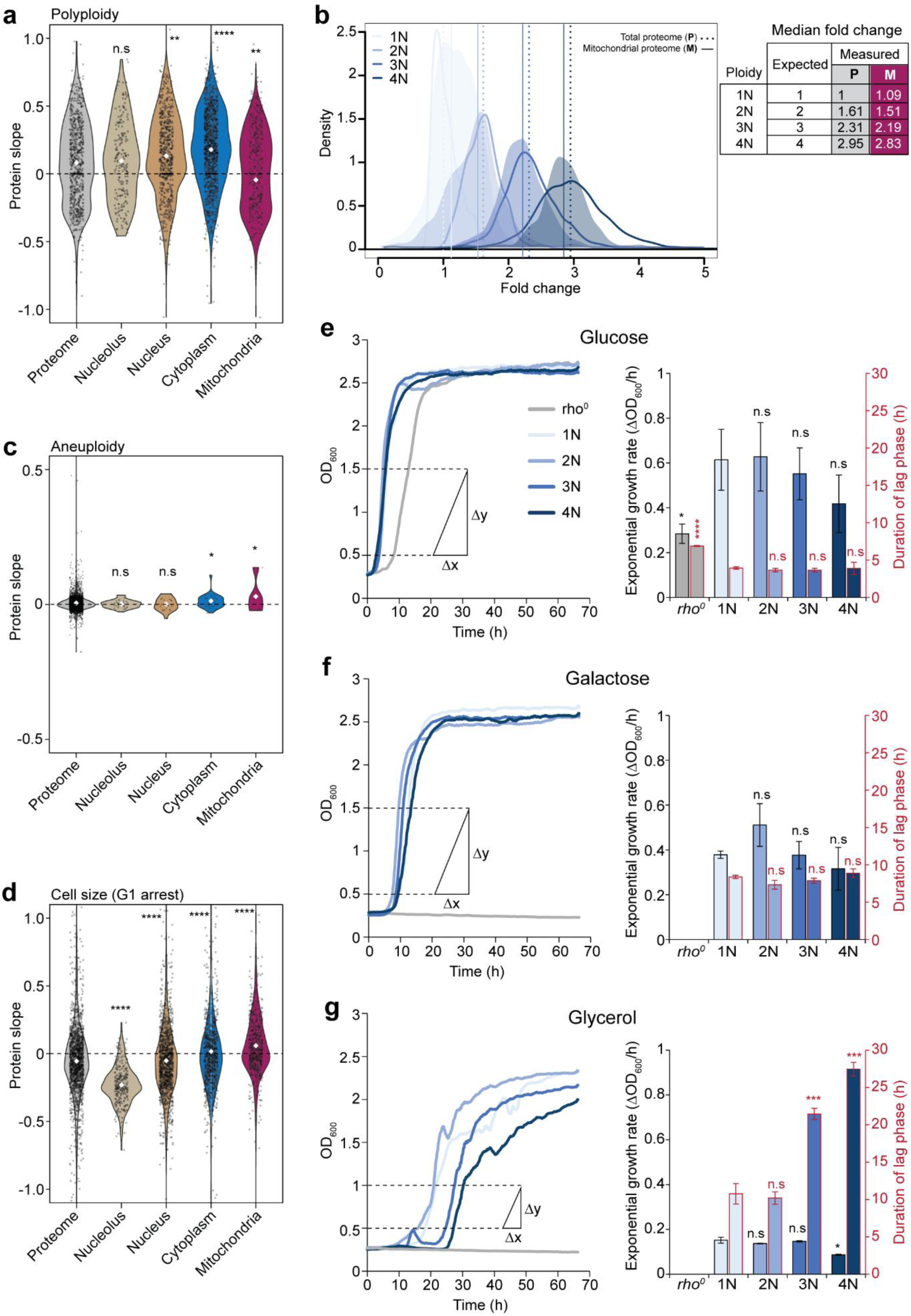
Polyploid budding yeasts exhibit poor scaling of mitochondrial proteins and growth defects under respiratory conditions. **a**, Influence of polyploidy on the scaling characteristics of proteins in budding yeast, calculated as protein slopes and sorted according to subcellular localization^12^. Proteins with a slope of 0 preserve their concentration independent of cell volume (i.e., proportional scaling), a slope up to 1 corresponds to an increase in protein concentration relative to increased cell volume (i.e., super-scaling) and a slope till -1 corresponds to a decrease in protein concentration relative to increased cell volume (i.e., sub-scaling). Each black dot represents a protein, and white dots represent mean value of the protein slope. *P*-values show statistical comparison to the global proteome and are derived from two-sided Wilcoxon’s ranks sum tests. * p < 0.05, ** p < 0.01, *** p < 0.001, **** p < 0.0001, n.s = not significant. **b,** Density plot comparing ploidy-dependent fold changes in expression of the global proteome and mitochondrial sub-proteome. Unfilled histograms represent distribution of the global proteome, with the median fold changes denoted by vertical dotted lines. Filled histograms represent distribution of the mitochondrial proteins, with the median fold changes denoted by solid vertical lines. The calculated median fold changes of protein expression are indicated in the table (right). All median fold changes were compared against the global protein expression in haploid strain, which was set to 1. Influence of **c,** aneuploidy^46^, and **d,** cell size^12^ on the scaling characteristics of proteins in budding yeast. **e-g,** Left: Growth curve of budding yeast ploidy series at 30°C in liquid media supplemented with **e,** glucose, **f,** galactose, or **g,** glycerol. Haploid *rho*^0^ yeast was included as control. OD600 range used for calculating slope of the exponential growth phase are shown. **e-g,** Right: Exponential growth rates and durations of the lag phase in **e,** glucose, **f,** galactose, and **g,** glycerol supplemented media. Data is shown as mean ± s.d. of three independent experiments. *P*-values represent two-tailed unpaired Student’s *t*-test. * p < 0.05, *** p < 0.001, **** p < 0.0001, n.s = not significant.

Yeast tetraploids are unstable and may assume aneuploid karyotypes^45^. To differentiate the influence of aneuploidy on protein scaling in polyploid yeasts, we calculated the protein slopes from previously published proteomics and cell volume data of isogenic aneuploid budding yeast series^46, 47^. Interestingly, the mean slopes of proteins localized to the various sub-compartments show a scaling behavior similar to the global proteome (mean slope; global proteome: 0, nucleolus: 0, nucleus: 0, cytoplasm: 0.01, mitochondria: 0.03) (Fig. 1c), thus indicating proportional scaling. To differentiate the influence of polyploidy from cell size on the scaling behavior of proteins in polyploid yeasts, we compared our results to similar analysis performed using budding yeast expressing the G1 cyclin, Cln1, under the control of a ß-estradiol-responsive promoter to generate cells of different sizes upon G1 arrest^12^. Here, a recalculation of protein slopes from the published proteomics dataset recapitulated previous results (mean slope; global proteome: -0.05, nucleolus: -0.23, nucleus: -0.05, cytoplasm: 0.02, mitochondria: 0.06) (Fig. 1d), confirming that the scaling of mitochondrial proteins in polyploid yeast is not controlled by cell size. Thus, the sub-scaling behavior of mitochondrial proteins in polyploid yeast is a ploidy-driven effect.

To further distinguish between the effect of ploidy or cell size on protein scaling behaviors across many different functional annotation groups, we performed a two-dimensional (2D) annotation enrichment analysis^48^ using the calculated protein slope values from our proteomics data on polyploid yeasts^11^ and published protein slope values from yeast of same ploidy but different sizes^12^. Strikingly, both mitochondrial (colored in magenta) and ribosome/translation-related (colored in blue) proteins sub-scale in response to polyploidy. In contrast, mitochondrial proteins super-scale, while ribosome/translation-related proteins sub-scale, in response to increasing cell size (Supplementary Fig. 1b, c). This observation is in line with our previous report that ribosome biogenesis and protein translation are dampened by increasing ploidy^11^, and suggests that this phenotype is driven by cell size scaling. Thus, we show that polyploidy remodels the cellular proteome by uncoupling the expression of mitochondrial proteins from increasing cell size, with components of the ETC, mt-DNA homeostasis, and antioxidant response pathways significantly repressed.

Next, we asked how the changes in the mitochondrial sub-proteome impact polyploid yeast proliferation under various growth conditions. Budding yeast cells can either undergo fermentation or respiration depending on the carbon source, which affects their growth and proliferation rate^49^. Using the isogenic budding yeast ploidy series^11^ (Supplementary Fig. 2a), we tested cell proliferation by measuring culture optical density at 600 nm (OD_600_) for 72 h at 30°C; as a control, we included an isogenic haploid *rho*^0^ yeast strain lacking the mitochondrial genome and therefore only grows under fermentative conditions. Yeasts of different ploidy grown in media supplemented with fermentative carbon (i.e., glucose and galactose) showed comparable growth, while growth of triploid and tetraploid strains was severely decreased on the non-fermentable carbon source glycerol (Fig. 1e-g). Particularly, both triploid and tetraploid yeast showed a significant growth delay in the lag phase - more than twice the time spent by haploid and diploid cells - while tetraploid yeast showed a statistically significant decrease in exponential growth rate in respiratory media (Fig. 1e-g). Additionally, both triploid and tetraploid yeast failed to grow to the same stationary phase density as haploid and diploid strains within the duration of OD_600_ measurement (Fig. 1e-g). Taken together, our results indicate that polyploid yeasts exhibit reduced mitochondrial proteome and growth defects under respiratory conditions, consistent with impaired mitochondrial respiratory functions.

### Polyploidy impairs scaling of mitochondrial content with increasing cell size

Mitochondrial network and mt-DNA content scale proportionally with cell size^21, 22^. Since cell size scales proportionally with ploidy in budding yeasts, we investigated scaling of the mitochondrial content in the yeast ploidy series. Firstly, we examined the surface area of the cell covered by mitochondrial network in two haploid and tetraploid yeast with different mating types. To do this, we transformed yeast with a genome-integrating mitochondria matrix-targeted mNeon green construct (i.e., Su9-mNeon green) and visualized fluorescence signal by confocal microscopy. While haploid (1N) yeast harbored an extended network of mitochondria, the network in the polyploid strains was sparsely distributed (Fig. 2a).

**Figure 2:**
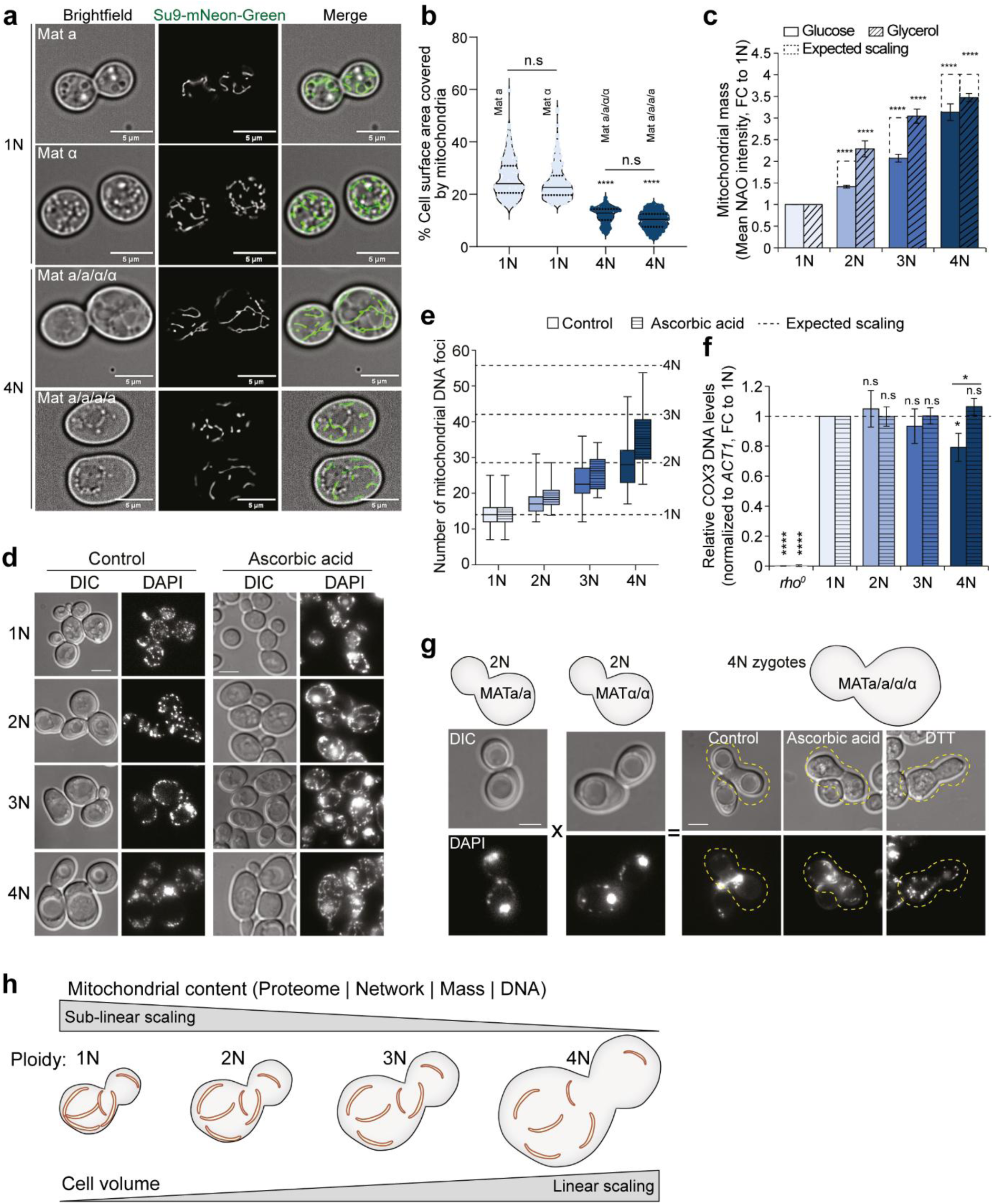
The mitochondrial content of polyploid yeast does not scale proportionally with cell size. **a**, Representative confocal images of the mitochondrial network in haploid (1N) and tetraploid (4N) yeasts of different mating types. Scale bar is 5 µm. **b,** Percentage of cell surface area covered by mitochondria from n = 187 (1N–Mat a), 269 (1N–Mat α), 140 (4N–Mat a/a/α/α), 245 (4N–Mat a/a/a/a) yeast cells in (**a**). Each dot represents one cell. The median (solid line) and interquartile range (dotted lines) of the distribution are shown. *P*-values represent non-parametric ANOVA (Kruskal-Wallis statistic = 630.6, p < 0.0001) followed by Dunn’s multiple comparisons test. Statistical comparisons were made against 1N – Mat a. **c,** Measurement of mitochondrial mass in yeast ploidy series. Data is shown as mean ± s.d. fold change to 1N, of three (glucose) and four (glycerol) independent experiments. The expected proportional scaling is indicated by dotted bars. *P*-values represent two-tailed unpaired Student’s *t*-test. **** p < 0.0001. **d,** Representative confocal images showing DNA staining in yeast ploidy series grown in media supplemented with or without ascorbic acid. Scale bar is 5 µm. **e,** Quantification of the number of mt-DNA foci in the cells from (**d**). Box plots show median foci number and interquartile range, while whiskers show minimum and maximum foci number extending from the lower and upper quartiles, respectively. Dotted horizontal lines indicate expected proportional scaling relative to 1N. **f,** Relative mt-DNA copy number in yeast ploidy series grown in media supplemented with or without ascorbic acid, determined by qPCR of the mitochondrially-encoded *COX3* normalized to nuclear-encoded *ACT1*. Haploid *rho^0^* cells were included as control. Data is shown as mean ± s.d. fold change to 1N, from three independent experiments. *P*-values represent two-tailed unpaired Student’s t-test. * p < 0.05, **** p < 0.0001, n.s = not significant. **g,** Representative confocal images showing the impact of ascorbic acid and DTT (i.e., reducing agents) on mt-DNA nucleoid abundance in freshly generated tetraploid zygotes. Scale bar is 5 µm. **h,** Schematic summarizing the scaling behavior of mitochondrial content (i.e., proteome, network morphology, mass and DNA abundance) with ploidy-dependent increase in cell volume.

Quantification of the percentage of cell surface area covered by mitochondria^50^ revealed a median value of approximately 25% in haploid yeast, which decreased to approximately 10% in tetraploid yeast (Fig. 2b). Next, we measured the mitochondrial mass by flow cytometry using normalized nonyl acridine orange (NAO) intensity, a dye that stains mitochondria independently of membrane potential. In comparison to the isogenic haploid, this revealed a 1.4-fold increase in mitochondrial mass in diploid, 2-fold in triploid and 3.1-fold in tetraploid yeasts upon growth in glucose-supplemented media, thereby indicating a sub-scaling of mitochondrial mass (Fig. 2c). Remarkably, the fold change in mitochondrial mass could be completely rescued in diploid and triploid, but not tetraploid, yeast upon growth in glycerol-supplemented media (Fig. 2c). Thus, mitochondrial network area and mass do not scale proportionally with ploidy-induced increase in cell size, but under severe metabolic demands such as during respiratory growth the scaling of the mitochondrial mass can be rescued, possibly due to stronger activation of stress response pathways.

Furthermore, we visualized and evaluated the number of mt-DNA nucleoids via 4′,6-diamidino-2-phenylindole (DAPI) staining and confocal microscopy. DAPI stains both nuclear and mitochondrial genetic material. To clearly distinguish nuclear DNA (nuc-DNA) staining from mt-DNA we either explored extra staining of mitochondria with Rhodamine dye to identify foci that colocalize with mitochondria or compared with parallel DAPI staining of *rho*^0^ cells lacking mt-DNA and hence showed exclusive nuc-DNA staining (Supplementary Fig. 2b, c). Mt-DNA packaging into nucleoids is impacted by oxidative stress^51, 52^. Given that polyploidy induces oxidative stress in fungal cells^53^, we additionally treated the cells with the reducing agent ascorbic acid prior to evaluation of mt-DNA nucleoids. This analysis revealed a ploidy-dependent increase in the number of mt-DNA nucleoids, which, however, did not scale proportionally with ploidy (Fig. 2d, e). Treatment with ascorbic acid further increased the number of mt-DNA foci (Fig. 2d, e), suggesting that polyploid yeasts exhibit oxidative stress that impacts mt-DNA homeostasis. Evaluation of the amount of mt-DNA by quantifying the levels of mitochondrially-encoded *COX3*, normalized to the nuclear-encoded *ACT1*, via quantitative PCR (qPCR) provided additional support for altered mt-DNA homeostasis. While *COX3* DNA levels were unchanged in diploid and triploid yeast, tetraploid yeast showed a significant 20% decrease in comparison to the haploid, which was rescued after treatment with ascorbic acid (Fig. 2f). Strikingly, the effect of oxidative stress on mt-DNA homeostasis appears to be an early event during polyploidization, as tetraploid zygotes obtained from freshly mated diploids showed low mt-DNA foci staining, which could be rescued by treatment with either ascorbic acid or dithiothreitol (DTT) (Fig. 2g). In summary, our results show that while polyploidy is associated with a proportional increase in cell size, mitochondrial proteome content, network coverage, mass and DNA amount do not scale proportionally (Fig. 2h) and may be linked to oxidative stress.

### Sub-linear scaling of mitochondrial content with ploidy is conserved in human cells

We asked whether the sub-linear scaling of mitochondrial content observed in polyploid yeast is conserved in mammalian cells. To this end, we sorted spontaneously arising near-tetraploid cells (labeled Spon) from a population of near-diploid human colorectal cancer HCT116 cell lines (i.e., 2N-Ctrl.)^7^. The status of p53, named guardian of the genome, influences the fate of tetraploid human cells and their ability to form tumors *in vivo*^4, 14, 54^. Therefore, to test the impact of p53 we additionally generated near-tetraploid cells by induction of cytokinesis failure using dihydrocytochalasin (DCD) treatment of p53-proficient (i.e., p53^+/+^-CF) and deficient (i.e., p53^-/-^-CF) HCT116 cells^5, 55^. The karyotype of the cells analyzed by flow cytometry confirmed near-tetraploid chromosomal content (Supplementary Fig. 3, Fig. 3a). The global proteome of the cells was measured via quantitative tandem mass tag (TMT)-labelled proteomics. Using this data, we calculated the scaling characteristics of each protein, sorted according to subcellular localization. Similarly to the polyploid yeasts, various proteins showed different scaling behaviors (Fig. 3b). The mean slopes of proteins localized to the different sub-compartments were significantly lower compared to the global proteome, with the scaling of mitochondrial proteins even more affected (mean slope; global proteome: 0.3, nucleolus: -0.02, nucleus: 0, cytoplasm: 0.02, mitochondria: -0.06) (Fig. 3b). Analysis of the log2-fold changes in abundance of individual mitochondrial proteins belonging to specific functional annotation groups revealed similar patterns to that observed in polyploid yeast (Supplementary Fig. 4).

**Figure 3:**
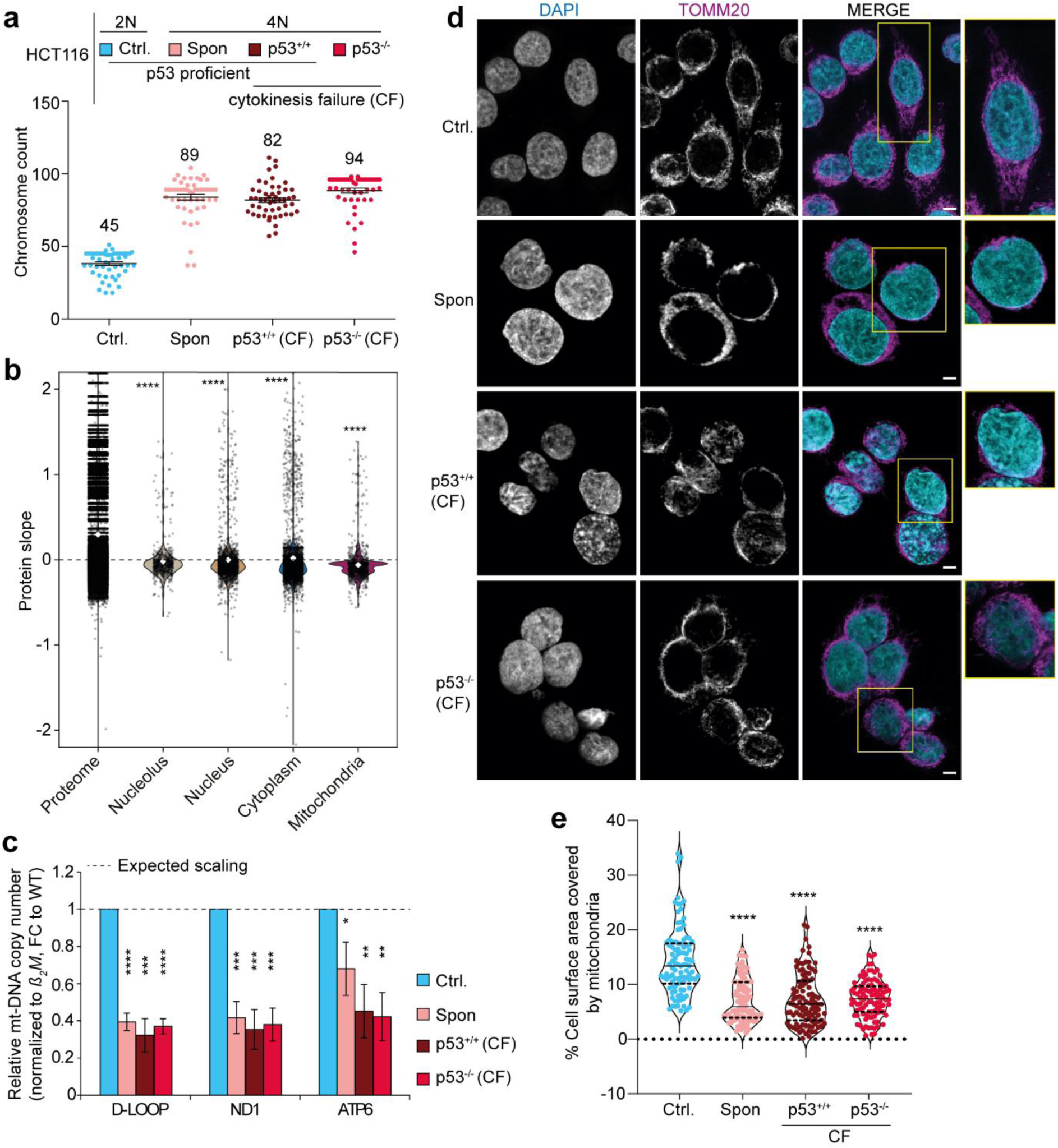
Ploidy-induced sub-scaling of mitochondrial content is conserved in human cells. **a**, Chromosome count of parental near-diploid human colorectal cancer HCT116 cells (i.e., 2N-Ctrl.), spontaneous near-tetraploids (4N) obtained by sorting from the 2N population (i.e., Spon) and derived near-tetraploids generated by induction of cytokinesis failure (CF) via dihydrocytochalasin D treatment in p53 proficient (p53^+/+^-CF) and deficient (p53^-/-^-CF) 2N cells. **b,** Scaling characteristics of proteins obtained by quantitative proteomics of the cells from (**a**), sorted according to subcellular localization. Protein slopes were calculated as described for Fig. 1a. Each black dot represents a protein, and white dots represent mean value of the protein slope. *P*-values show statistical comparison to the global proteome and are derived from two-sided Wilcoxon’s ranks sum tests. **** p < 0.0001. **c,** Relative mt-DNA copy number determined by qPCR of the non-coding mt-DNA *D-LOOP*, respiratory chain complex 1 subunit *ND1*, and respiratory chain complex 5 subunit *ATP6* normalized to nuclear-encoded ß2-microglobulin (*ß2M*). Data is shown as mean ± s.d. fold change to the parental diploid (2N) from three independent experiments. *P*-values represent two-tailed unpaired Student’s t-test. * p < 0.05, ** p < 0.01, *** p < 0.001, **** p < 0.0001. **d,** Representative confocal images of the mitochondrial network in the different cell lines in (**a**), visualized by immunofluorescence of the translocase of outer mitochondrial membrane TOMM20. DAPI is used as a nuclear counterstain. Scale bar is 10 µm. **e,** Percentage of cell surface area covered by mitochondria from n = 100 cells from (**d**). Each dot represents one cell. The median (solid line) and interquartile range (dotted lines) of the distribution are shown. *P*-values represent non-parametric ANOVA (Kruskal-Wallis statistic = 104.1, p < 0.0001) followed by Dunn’s multiple comparisons test. Statistical comparisons were made against 2N-Ctrl. cells.

Next, we quantified the mt-DNA copy number through qPCR, using primers specific to the non-coding mt-DNA *D-LOOP* and coding sequences of respiratory chain complex I (i.e., *ND1*) and V (i.e., *ATP6*) proteins. The mt-DNA levels were normalized to that of the nuclear-encoded ß_2_-microglobulin (*ß_2_M*) gene. This revealed nearly 60% decrease in mt-DNA levels in tetraploid human cells (Fig. 3c). Furthermore, we analyzed the mitochondrial network in diploid and tetraploid human cells by immunostaining of the outer membrane translocase TOMM20 and, visualizing by confocal microscopy, observed reduced network morphology in tetraploid cells compared to parental diploids (Fig. 3d). Quantification of the percentage of cell surface area covered by mitochondria^56, 57^ showed a median decrease of approximately 50% in tetraploid compared to diploid cells (Fig. 3e). Intriguingly, the observed mitochondrial phenotypes were independent of p53 status and mode of tetraploidization. Taken together, the sub-linear scaling of mitochondrial content to ploidy is conserved in mammalian cells.

### Whole genome duplication causes defects in mitochondrial functions

We reasoned that the altered mitochondrial contents upon polyploidization would affect mitochondrial functions. To this end, we investigated several phenotypes that indicate mitochondrial health using the yeast ploidy series. Firstly, we isolated mitochondria from cells grown in media supplemented with galactose and examined *in organello* translation capacity via incorporation of radio-labelled methionine (^35^S-Methionine) into newly synthesized proteins. Samples were resolved by SDS-PAGE and visualized by autoradiography. The yeast mitochondrial genome encodes eight proteins; seven are essential subunits of ETC complexes and one is a structural ribosomal protein^58^. We included *rho*^0^ cells as a control. Strikingly, we observed small, but significant decrease in mitochondrial translation capacity in tetraploid cells at the time point 15 min (Fig. 4a, b), suggesting that tetraploidy limits translation of the mitochondrial genome.

**Figure 4:**
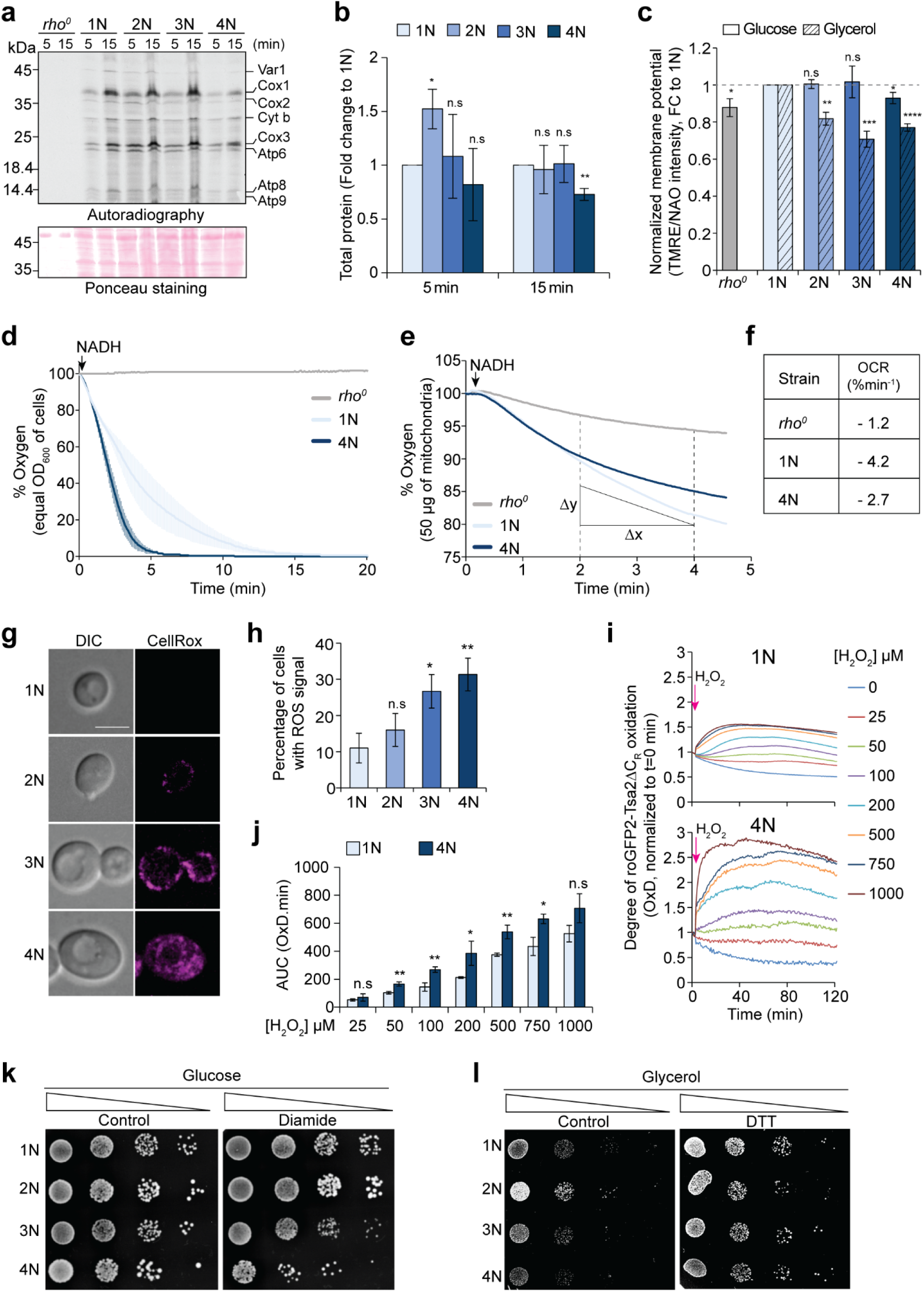
Polyploidy causes functional mitochondrial defects and increased oxidative stress. **a**, Representative autoradiograph of radiolabeled translation products from mitochondria isolated from the yeast ploidy series. Mitochondria from *rho^0^* cells were included as control. **b,** Quantification of total protein amount from (**a**), normalized to ponceau staining. Data is shown as mean ± s.e.m fold change to 1N, from three independent experiments. *P*-values represent two-tailed unpaired Student’s *t*-test. * p < 0.05, ** p < 0.01, n.s = not significant. **c,** Measurement of mitochondrial membrane potential in yeast ploidy series. Data is shown as mean ± s.d. fold change to 1N, from three independent experiments. *P*-values represent two-tailed unpaired Student’s *t*-test. * p < 0.05, ** p < 0.01, *** p < 0.001, **** p < 0.0001, n.s = not significant. **d,** Measurement of NADH-induced oxygen consumption of mitochondria isolated from equal OD600 of yeast cells. **e,** Measurement of NADH-induced oxygen consumption of 50 µg mitochondria isolated from yeast cells. Duration used for calculating slope of oxygen consumption is indicated. **f,** Oxygen consumption rate (OCR) determined from the slopes in (**e**). **g,** Representative confocal images of reactive oxygen species (ROS) detection in yeast ploidy series grown in glycerol supplemented media, using CellROX DeepRed. Scale bar is 5 µm. **h,** Quantification of the percentage of cells with positive CellROX staining from (**g**). **i,** Degree of roGFP2 oxidation in haploid and tetraploid yeast in response to different concentrations of hydrogen peroxide (H2O2). Data is normalized to the start timepoint of measurement prior to H2O2 addition. **j,** Quantification of the area under the curve for the different concentrations of H2O2 relative to the untreated sample in (**i**); depicting H2O2 scavenging capacity. Sensitivity of yeast ploidy series to growth on **k,** glucose-supplemented agar treated with or without diamide (i.e., oxidizing agent) and **l,** glycerol-supplemented agar treated with or without DTT (i.e., reducing agent).

Secondly, we evaluated mitochondria membrane potential by staining yeast cells with the membrane-permeable fluorescent dye tetramethylrhodamine ethyl ester perchlorate (TMRE), whose accumulation in mitochondria is membrane potential-dependent, and measured dye intensity by flow cytometry. The data was normalized to the fluorescent intensity of NAO, which stains all mitochondria. Only tetraploid yeast exhibited significantly decreased membrane potential upon growth in fermentative media, similar to *rho*^0^ cells (Fig. 4c). However, the membrane potential was significantly decreased in respiratory media with increasing ploidy, suggesting that polyploidy exacerbates defects in mitochondria membrane potential under increased metabolic demands (Fig. 4c).

Mitochondria membrane potential is linked to respiratory chain functions. Therefore, we analyzed and compared the respiratory capacity of mitochondria isolated from haploid and tetraploid yeast grown in media supplemented with galactose via NADH-driven oxygen consumption *in vitro*. Here, we used two approaches: (i) equal aliquot of isolated mitochondria from an equal OD_600_ of yeast cells was used for the oxygen consumption assay without adjusting the mitochondrial amount, and (ii) the mitochondrial concentration was measured via Bradford assay and equal amounts were used for the assay (i.e., 50 µg). This allowed us to distinguish between oxygen consumption capacity influenced by total mitochondria amount and effective mitochondria oxygen consumption capacity per strain. In both analyses, we included *rho*^0^ cells as control. In the first scenario, mitochondria from tetraploid yeast consumed oxygen at a faster rate than haploid yeast, consistent with higher total mitochondrial amount per OD_600_ units of cells although the mitochondrial mass does not scale proportionally with cell size (Fig. 4d). In the second scenario, equal amount of mitochondria from tetraploid cells showed nearly 30% decrease in oxygen consumption rate (OCR) compared to mitochondria from haploids, suggesting that mitochondria in haploid cells are more respiration-competent than tetraploids (Fig. 4e, f). Thus, mitochondrial membrane potential and respiratory capacity are significantly reduced in tetraploid cells.

Further, we asked whether the mitochondrial respiratory defect is associated with altered redox homeostasis, and thus oxidative stress. To this end, we examined the levels of ROS by staining yeast cells grown in glycerol-supplemented media with CellROX Deep Red reagent, a chemical dye that fluorescently labels ROS species, and quantified the proportion of cells with CellROX staining via microscopy. We observed significantly increased percentage of CellROX-stained cells in triploid and tetraploid yeast (Fig. 4g, h). To further explore the redox changes, we transformed haploid and tetraploid yeast with the genetically encoded roGFP2-Tsa2ΔC_R_ biosensor to detect hydrogen peroxide (H_2_O_2_)^59, 60^. The degree of roGFP2 oxidation (OxD), in yeast grown in glucose-supplemented media, under basal unperturbed state and in response to different concentrations of H_2_O_2_ was analyzed to assess the cellular ROS scavenging capacity. The ROS scavenging capacity was evaluated as the area under the curve (AUC) between a H_2_O_2_-treated and untreated condition; a higher AUC is indicative of reduced scavenging capacity^61, 62^. Tetraploid yeasts were more sensitive than haploid yeasts to the different concentrations of H_2_O_2_ and showed poorer scavenging capacity (Fig. 4i, j). The sensitivity to ROS was further supported by the observation that triploid and tetraploid yeasts showed decreased growth on fermentative media supplemented with the oxidizing agent diamide (Fig. 4k). In contrast, growth defects of triploid and tetraploid yeast on respiratory media could be rescued by supplementation with the reducing agent DTT (Fig. 4l). Together, these results reveal profound changes in mitochondrial metabolism and redox homeostasis in polyploid budding yeasts.

Mitochondrial dysfunction has also been linked to cytosolic protein aggregation stress arising from gene dosage imbalance in aneuploid cells^63, 64^. Since polyploid cells are inherently unstable, we asked if protein aggregation potentially contributes to the observed mitochondrial defects. Hence, we investigated the formation of cytosolic protein aggregates by transforming haploid and tetraploid yeasts with a plasmid construct of Hsp104-GFP under the expression of a constitutive promoter, and visualized Hsp104 foci by confocal microscopy. Hsp104 is a molecular chaperone that localizes to protein aggregates, and formation of Hsp104 foci is indicative of cytosolic protein aggregation^65^. As a control, we subjected the cells to a heat shock at 37°C for 2h to induce protein aggregation. In both strains we observed either diffused GFP signal throughout the cytoplasm or GFP signals in puncta (Supplementary Fig. 5a). Under normal unperturbed conditions, both haploid and tetraploid yeast harbored cells with Hsp104 puncta, consistent with basal cytosolic proteostasis, which were increased by heat shock (Supplementary Fig. 5a). Quantification of the proportion of cells harboring either diffused or punctate GFP revealed no significant differences between the haploid and tetraploid cells (Supplementary Fig. 5b). This observation was consistent with the abundance of molecular chaperones when compared against the global proteome in the ploidy series, which showed no significant differences (Supplementary Fig. 5c). Thus, the mitochondrial defects in polyploid yeast, unlike in aneuploid cells, cannot be attributed to cytosolic protein aggregation stress.

### The mitochondrial retrograde signaling pathway rescues mitochondrial mass in polyploid yeast

Our observation that mitochondrial mass scaled nearly linearly with ploidy under respiratory conditions, in contrast to the poor scaling observed under fermentative growth (Fig. 2c), was unexpected and prompted us to investigate the underlying mechanism. In budding yeast, mitochondrial mass positively correlates with mitochondrial metabolism during respiratory growth or biosynthetic demand, thereby linking changing mitochondrial mass to changing organelle function^66^. Mitochondrial metabolism is driven by the heme activator protein complex (HAP) system under normal physiological conditions (e.g., during diauxic shift), and by the retrograde (RTG) system under both physiological and dysfunctional states (Fig. 5a). Both pathways activate transcription of diverse but also similar genes, including components of the TCA cycle^36, 67, 68^. We previously observed that the main effector of the RTG pathway, Rtg2, is among the top 25 most abundant proteins across all polyploid yeast strains and its levels are significantly stabilized in a ploidy-dependent manner, as revealed by abundance analysis upon translation inhibition with cycloheximide^11^. Our analysis of transcriptomics data showed no significant changes in the transcript levels of the HAP pathway genes in polyploid yeast. By proteomics, the Hap2 protein was significantly reduced in abundance with increasing ploidy; no other HAP pathway proteins were quantified (Fig. 5b). Thus, we found no evidence for robust activation of the HAP pathway in response to ploidy changes; at best we observed pathway suppression. In contrast, although the expression of many RTG pathway candidates is mostly unchanged at the transcript level, their protein abundance, including that of TCA cycle components, is significantly increased in polyploid yeast (Supplementary Fig. 1a, Fig. 5b). Thus, the RTG system is activated in response to polyploidization in budding yeast, possibly due to the increased mitochondrial defects.

**Figure 5:**
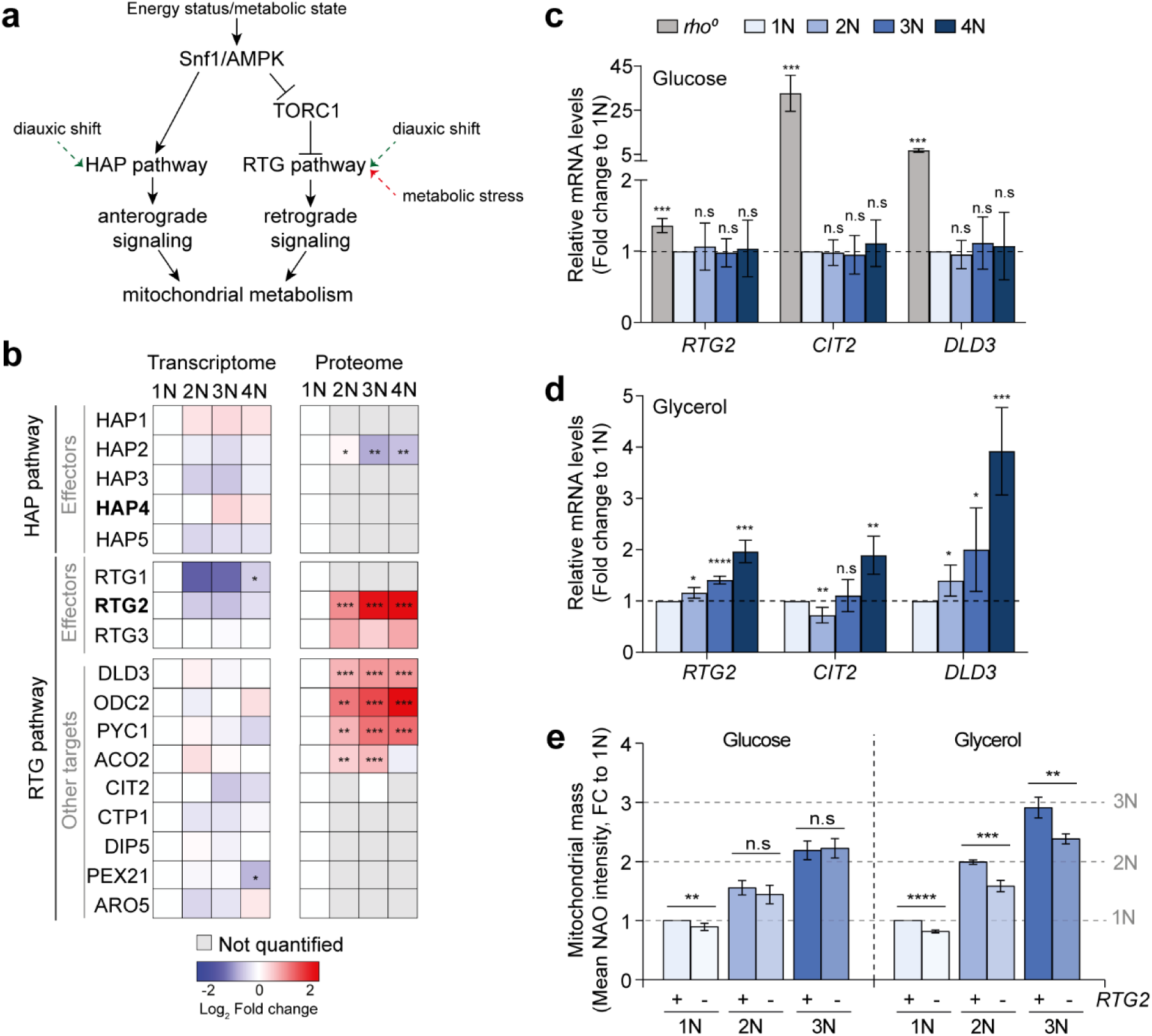
The RTG pathway regulates mitochondrial homeostasis in polyploid yeast. **a**, Schematic depiction of the two main pathways regulating mitochondrial metabolism in budding yeast under normal physiological conditions (HAP pathway, anterograde signaling) and metabolic stress (RTG pathway, retrograde signaling). **b,** Heatmap showing Log2 fold change in expression, relative to the haploid (1N) strain, of pathway genes from the transcriptome and proteome of the yeast ploidy series. Grey boxes indicate values not statistically quantified. * q < 0.05, ** q < 0.01, *** q < 0.001. Quantitative PCR of selected RTG pathway genes from the yeast ploidy series grown in either **c,** glucose or **d,** glycerol supplemented media, normalized to *ACT1*. Haploid *rho^0^*cells were included as control. Data is shown as mean ± s.d. fold change to 1N, from three independent experiments. *P*-values represent two-tailed unpaired Student’s t-test. * p < 0.05, **** p < 0.0001, n.s = not significant. **e,** Measurement of mitochondrial mass in budding yeast possessing or lacking *RTG2*. Data is shown as mean ± s.d. fold change to 1N, of five (glucose) and four (glycerol) independent experiments. The expected proportional scaling is indicated with dotted lines. *P*-values represent two-tailed unpaired Student’s *t*-test. ** p < 0.01, *** p < 0.001, **** p < 0.0001, n.s = not significant.

Analysis of the expression levels of RTG pathway genes by RT-qPCR revealed no significant changes under fermentative growth, but significantly increased expression under respiratory growth (Fig. 5c, d). To further examine the relevance of the RTG pathway for polyploid yeast survival, we investigated the impact of *RTG2* loss. To this end, we obtained a library of haploid *rtg2Δ* strains of opposite mating types and generated diploid *rtg2Δ* strains by replacing *RTG2* with antibiotic-resistant cassettes in parental diploids of different mating types^11^. Haploids and diploids with and without *RTG2* were mated with each other to obtain triploid and tetraploid progenies. Surprisingly, while loss of *RTG2* did not affect yeast mating phenotype and zygote formation (Supplementary Fig. 6, white circles), we were unable to generate viable tetraploid *rtg2Δ* progenies, suggesting that *RTG2* is required for efficient establishment or survival of tetraploid cells.

Next, we evaluated the consequence of *RTG2* loss on mitochondrial mass via flow cytometry. In haploid yeast grown under fermentative conditions loss of *RTG2* significantly decreased mitochondrial mass by approximately 10%, suggesting that the RTG pathway contributes to mitochondria biogenesis even in the absence of metabolic stress (Fig. 5e). In contrast, the mitochondrial mass of the haploid *rtg2Δ* strain decreased by approximately 20% under respiratory growth (Fig. 5e). While mitochondrial mass of diploid and triploid *rtg2Δ* yeasts was not significantly altered under fermentative growth, it was significantly reduced by approximately 20% in both strains under respiratory growth, in comparison to the *RTG2*-proficient parental (Fig. 5e). Taken together, our data suggests that Rtg2 is important for maintaining mitochondrial mass, hence metabolism, in polyploid yeast and its loss may be incompatible with viability in tetraploid yeast or yeasts with higher ploidies.

### Stress-activated kinase Hog1 and TORC1 coregulate mitochondria retrograde signaling in yeast

What exactly is the upstream trigger of the RTG pathway in polyploid cells? The RTG pathway is governed by mitogen and stress-activated protein kinases (MAPK and SAPK), which regulate pathway activation via its phosphorylation^41^. Our proteomics analysis showed that the levels of cell integrity and stress-activated MAPKs, Slt2 and Hog1, respectively, were significantly increased in polyploid yeasts (Fig. 6a). Notably, the Hog1 (high osmolarity glycerol) SAPK, which directly regulates the RTG pathway^41, 69^, is activated via dual phosphorylation of the highly conserved threonine 174 (Thr174) and tyrosine 176 (Tyr176) residues^70, 71^ or by oxidation of conserved thiols at Cys156, Cys161 and Cys205^41, 72–74^ in response to different stressors. Hog1 protein levels were slightly, but significantly, increased in polyploid yeast under fermentative growth and significantly increased in a ploidy-dependent manner under respiratory conditions (Fig. 6b, c). However, while Hog1 phosphorylation could be strongly induced upon osmotic shock with NaCl treatment in haploid yeast, it was barely observed in polyploid yeast grown under both fermentative and respiratory conditions (Fig. 6d, e), raising the possibility that Hog1 abundance, rather than canonical phosphorylation, contributes to polyploidy-induced retrograde signaling.

**Figure 6:**
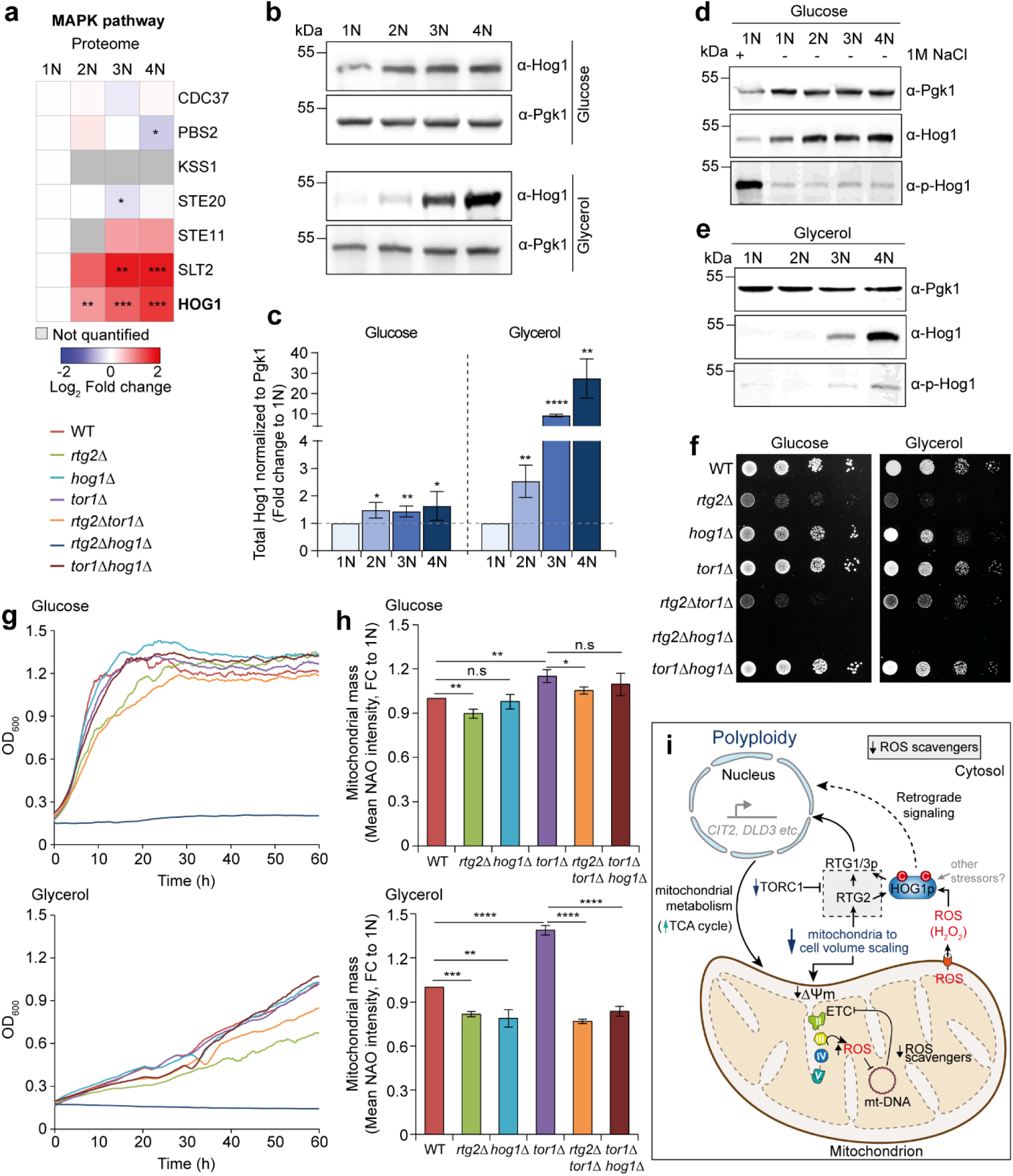
TORC1 and HOG1 coregulate the RTG pathway to control mitochondrial metabolism in budding yeast. **a**, Heatmap showing Log2 fold change in expression, relative to the haploid (1N) strain, of MAPK pathway proteins from the proteome of the yeast ploidy series. Grey boxes indicate values not statistically quantified. * q < 0.05, ** q < 0.01, *** q < 0.001, *** q < 0.0001. **b,** Representative immunoblot of Hog1 protein levels in the yeast ploidy series. Pgk1 was used as loading control. **c,** Quantification of the Hog1 protein levels from (**b**). Data is shown as mean ± s.d. fold change to 1N, from three independent experiments. *P*-values represent two-tailed unpaired Student’s *t*-test. * p < 0.05, ** p < 0.01, **** p < 0.0001. Analysis of Hog1 activation by phosphorylation at Thr174/Tyr176 in yeast ploidy series grown in **d,** glucose and **e,** glycerol media, via immunoblot. Haploid yeast treated with 1M NaCl to induce osmotic shock was included as positive control for Hog1 phosphorylation. Pgk1 was used as loading control. **f,** Yeast growth on agar by serial dilution. **g,** Yeast growth curve in liquid media supplemented with glucose or glycerol. Pre-cultures were diluted to 0.1 OD600 prior to growth on agar plates (**f**) or liquid media (**g**). **h,** Measurement of mitochondrial mass in WT budding yeast and gene deletion variants. Data is shown as mean ± s.d. fold change to three independent experiments. *P*-values represent two-tailed unpaired Student’s *t*-test. * p < 0.05, ** p < 0.01, *** p < 0.001, **** p < 0.0001, n.s = not significant. **i,** Schematic of the proposed mechanism of mitochondrial retrograde signaling in response to polyploidy-induced sub-linear scaling of mitochondrial content to cell volume.

Our previous work revealed that the activity of the target of rapamycin (TOR) complex TORC1 was significantly decreased in polyploid compared to haploid yeast^11^. Whereas Hog1 is a positive regulator, TORC1 is a negative regulator of the RTG pathway acting either upstream of Rtg2p or between Rtg2p and Rtg1p/3p^42, 75, 76^. Therefore, we investigated the interplay of Rtg2, Hog1 and Tor1 on yeast survival/proliferation and mitochondrial metabolism via gene deletion and phenotypic analyses. Loss of *RTG2* partially impaired yeast proliferation under fermentative growth, which was exacerbated under respiratory conditions (Fig. 6f, g). Loss of *HOG1* did not affect yeast proliferation under fermentative conditions, but the *hog1Δ* strain showed reduced growth under respiratory conditions (Fig. 6f, g). In contrast, yeast with *TOR1* deletion (the only subunit of TORC1 whose deletion is compatible with cell viability in yeast) showed partial growth advantage under fermentative conditions and its growth was not altered under respiratory conditions (Fig. 6f, g). Loss of *RTG2* in a *tor1Δ* strain recapitulated the growth defects under both fermentative and respiratory conditions (Fig. 6f, g). While deletion of *TOR1* in the *hog1Δ* strain showed no significant changes under fermentative growth, it moderately improved respiratory growth in comparison to the *hog1Δ* strain alone. Strikingly, however, yeast with combined loss of *RTG2* and *HOG1* were viable but did not grow beyond an initial pre-culture phase (Fig. 6f, g).

Next, we analyzed the impact of *RTG2*, *HOG1* and *TOR1* deletion on mitochondrial metabolism by measuring the mitochondrial mass. Loss of *RTG2* significantly reduced mitochondrial mass in both WT, by 10% and 20%, and *tor1Δ* strains, by 10% and 45%, under fermentative and respiratory conditions, respectively, while loss of *HOG1* achieved similar effect only under respiratory growth (Fig. 6h). In contrast, loss of *TOR1* was beneficial for mitochondrial mass under both fermentative and respiratory growth (Fig. 6h). Thus, Rtg2 is important for mitochondrial metabolism, which correlates with yeast proliferation, under both fermentative and respiratory conditions, while Hog1 is dispensable for fermentative growth.

Taken together, our results suggest that the suppression of TORC1 activity, stabilization of Rtg2 protein levels^11^ and Hog1 protein abundance in polyploid yeast may constitute an adaptive response to ploidy-induced mitochondrial defects, which sustains mitochondrial metabolism (e.g., via the TCA cycle) through retrograde signaling to ensure yeast survival/proliferation under increased metabolic demands despite chronic mitochondrial respiratory defects. We propose that the poor scaling of mitochondrial content to polyploidy-induced increase in cell volume instigates mitochondrial respiratory defects leading to increased ROS production and oxidative stress. Cellular ROS in the form of H_2_O_2_ mediates activation of the Hog1 SAPK, thereby promoting mitochondrial metabolism through retrograde signaling. Finally, Tor1 negatively regulates the signaling mechanism (Fig. 6i).

## Discussion

Organellar homeostasis is critical for healthy cellular and organismal physiology. Polyploidy, a phenomenon characterized by increased number of complete chromosome sets, is usually associated with a proportional increase in nuclear and cellular volume, elevated genome instability and growth defects across diverse taxa. At the same time, polyploidy is frequent in nature, where it confers evolutionary advantages for cell and organismal resilience under stress^53, 77–80^. Furthermore, nearly 40% of human cancers underwent at least one round of whole genome duplication^81^. Yet, the consequences of polyploidy in evolution and cancer and how it facilitates adaptation and tumor growth remain incompletely understood. By quantitative proteomics analysis of an isogenic series of budding yeast *S. cerevisiae*, from haploid to tetraploid, we previously showed that polyploidy is associated with sub-linear scaling of the cellular proteome due to decreased ribosome biogenesis and protein translation^11^. Here, we show that while proteins localized to the nucleolus, nucleus and cytoplasm scale with ploidy, mitochondrial proteins exhibit a strong sub-scaling behavior (Fig. 1a). Strikingly, components of the respiratory chain, proteins involved in mt-DNA maintenance, and redox homeostasis are significantly depleted in polyploid yeasts. In contrast, enzymes of the TCA cycle are significantly increased in response to polyploidization (Supplementary Fig. 1a). This poor scaling of mitochondrial proteins is a polyploidy-dependent phenotype, not driven by aneuploidy or cell size (Supplementary Fig. 1b, c, Fig. 1b-d), and conserved in near-tetraploid cells derived from HCT116, a human colorectal cancer cell line (Supplementary Fig. 4, Fig. 3a, b). Understanding how mitochondrial proteome sub-scaling upon polyploidization impacts the cell’s metabolic homeostasis and survival under diverse growth conditions is key to unravelling the link between polyploidy and metabolic reprogramming in cancer.

What is/are the cause(s) of mitochondrial scaling and respiratory defects in polyploid cells? We reasoned that mitochondria biogenesis, e.g., protein import, may be impaired. In fact, the changes we observed in the mitochondrial proteome of polyploid cells (Supplementary Fig. 1a) are surprisingly similar to those observed in cells in which the import of mitochondrial proteins was competitively inhibited by use of clogger constructs that blocked protein translocation into the organelle^82^. Reduced mitochondrial protein import efficiency is linked with suppression or inactivation of the HAP transcription complex to downregulate expression of OXPHOS components, in an adaptive response that prevents accumulation of unimported precursor proteins^82^. Our observation of reduced abundance of Hap2 and OXPHOS components in polyploid cells is consistent with this finding (Supplementary Fig. 1a, Fig. 5b). However, it should be noted that unlike the clogger model, which represents acute stress, polyploidy is chronic. An alternative hypothesis is that dilution of the mitochondrial proteome potentially alters the protein to lipid composition of mitochondrial membranes, thereby affecting membrane-related functions such as cristae formation, ETC assembly and respiration. We are optimistic that future investigations will shed more light on these propositions.

Furthermore, the rescue of mt-DNA levels in tetraploid yeast via antioxidant agents suggests a redox-mediated regulation of mt-DNA homeostasis in response to polyploidy. This is also reflected in proliferation capacity: while polyploid yeasts generally show comparable growth to haploids under fermentative conditions, they are more sensitive to growth in the presence of oxidizing agents and the growth defects under respiratory conditions are rescued through reducing agents (Fig. 1e-g, 4k-l). Moreover, although the mitochondria mass of polyploid yeast does not scale linearly with ploidy, the scaling defect is almost completely rescued under respiratory growth (Fig. 2c); a condition that leads to increased ROS production (Fig. 4g). Together, these observations highlight a connection between redox homeostasis, mitochondria biogenesis and cell survival. Similar increased ROS production was previously observed in tetraploid *Candida albicans*, where it contributes to concerted ploidy reduction^53^. It should be noted that despite its cytotoxic effects, mitochondria-derived ROS also plays important physiological signaling functions within the cell, mainly via thiol redox switches ^83–85^. How redox sensing may be linked to mitochondria biogenesis is a subject for future research.

Mitochondrial defects were reported in response to aneuploidy in *Drosophila* and human cells, but attributed to cytosolic protein aggregation stress^63, 64, 86^. However, cytosolic proteostasis defects are not heightened in response to polyploidization (Supplementary Fig. 5) suggesting that the consequences of ploidy changes for mitochondrial homeostasis in aneuploid and polyploid cells are manifested via two distinct mechanisms; accumulation of cytosolic protein aggregates that sequester mitochondrial preproteins in aneuploidy, and sub-scaling of mitochondrial content to increased cell volume in polyploidy, leading to mitochondrial dysfunctions and metabolic stress.

What then is the consequence of mitochondrial scaling and respiratory defects in polyploid yeast? In contrast to HAP, we find that the RTG pathway is activated in polyploid yeast (Supplementary Fig. 1a, Fig. 5b-d). In budding yeast with compromised mitochondrial respiratory functions, the increased expression of genes involved in the early steps of the TCA cycle – i.e., *CIT1* (encoding a mitochondrial isoform of citrate synthase), *ACO1* (encoding aconitase), and *IDH1/2* (encoding NAD^+^-dependent isocitrate dehydrogenase) – as well as glyoxylate cycle genes (e.g., *CIT2*, *DLD3*, *PYC1*) that supply acetyl-CoA and oxaloacetate to the TCA cycle, is consistent with retrograde signaling^31, 36^. This expression changes drives metabolic readjustment by permitting mitochondrial metabolism via the TCA cycle, enabling yeast to bypass mitochondrial respiratory deficiency for sustained energy production, amino acid and nucleotide biosynthesis needed for proliferation^31^. Disease mutations in *MT-ATP6* (ETC complex V component encoded from mt-DNA), which causes chronic ATP synthase dysfunction and respiration defects that underlie the NARP syndrome, triggers increased TCA cycle activity in yeast and Drosophila as metabolic compensation to sustain energy availability^87, 88^. In yeast, such compensation is driven by the retrograde pathway through changes in NAD^+^/NADH metabolism^88^. Our observed increase in abundance of the NADH dehydrogenases, Nde1 and Ndi1, in polypoid yeast potentially reflects changes in NAD^+^/NADH metabolism. Interestingly, while mitochondrial mass increased and scaled nearly linearly with ploidy under respiratory growth (Fig. 2c), the mitochondrial membrane potential significantly decreased (Fig. 4c), suggesting that the mitochondrial metabolic capacity of a cell may be compensated despite ETC/OXPHOS deficiency. Future metabolomics analyses will reveal how polyploidy-induced retrograde signaling drives metabolic adaptation for yeast survival. Many cancers exhibit a metabolic shift from OXPHOS to glycolysis via the Warburg effect, thereby ensuring fast energy production via glycolysis under aerobic conditions and providing essential metabolic intermediates needed for cell growth, survival and proliferation^89^. While diverse mechanisms have been proposed to explain this phenomenon, the influence of polyploidy described here could add a missing piece to the puzzle.

How may the polyploidy-induced retrograde response be regulated? Although canonical retrograde signaling occurs via transcriptional activation^90^, we find, unexpectedly, that several components of the RTG pathway, including TCA cycle components, accumulate at the protein level without corresponding changes in transcript abundance (except when respiration is induced), indicating that the pathway activation can occur independently of transcription induction (Fig. 5b-d). It is plausible that the transcriptional response occurred early post-polyploidization, as with the mitochondrial defects (Fig. 2g), and abated when the retrograde pathway protein levels stabilized. While the underlying mechanism remains to be determined, this finding is consistent with the notion that post-transcriptional regulation, such as altered protein stability, strongly contributes to the cellular response to chronic or persistent stress. We propose that such unconventional regulation may provide a rapid and economical mechanism for modulating retrograde signaling in response to mild but chronic mitochondrial dysfunction.

The essentiality of the retrograde response for polyploid yeast survival and proliferation is reflected in the observation that loss of *RTG2* reduces mitochondrial mass, thus metabolism, in yeasts of any ploidy and appears incompatible with survival of tetraploids or yeast of higher ploidies (Supplementary Fig. 6, Fig. 5e). Hog1 abundance may serve an additional role under increased metabolic demands, while loss of both *RTG2* and *HOG1* may be detrimental to the retrograde response and yeast proliferation^69^ (Fig. 6f-h). The RTG and HOG pathways are activated in response to a variety of stressors, including oxidative stress. Polyploid yeasts increase ROS production, which we hypothesize mediates the coupling of mitochondrial dysfunction to the retrograde response. How this sensing mechanism may be achieved partially lies in the regulation of Hog1. Hog1 is traditionally activated via phosphorylation at conserved threonine and tyrosine residues during osmotic stress^70, 71^. However, it also possesses conserved cysteines whose thiols can be modified to regulate Hog1 activity upon exposure to ROS^72, 74^. Indeed, redox regulation of Hog1 and not its phosphorylation was required for the response to nitrosative stress in *C. albicans*^73^. In this strain, a master regulator of Hog1, i.e., Ssk1, maintains a basal level of Hog1 phosphorylation to facilitate the response to oxidative but not osmotic stress^91^. Remarkably, Hog1 phosphorylation was barely induced in polyploid yeast despite increase in protein abundance (Fig. 6a-e), compelling us to propose that thiol redox modification of conserved cysteine residues^73, 74, 92^ may be crucial for Hog1 activation to drive retrograde signaling in polyploid yeast (Fig. 6h). Furthermore, oxidative stress mediated activation of Hog1 can also be achieved through redox-sensitive inactivation of protein tyrosine phosphatases that negatively regulate Hog1 (e.g., Ptp2 and Ptp3)^91, 93^. Given that we did not observe any increase in Hog1 transcript levels in polyploid yeasts^11^, increased abundance of the Hog1 protein may reflect sustained pathway activation or post-transcriptional regulation rather than transcriptional induction. Such regulation may also occur to safeguard genome integrity in response to polyploidy-induced chronic oxidative stress^94^.

A counterintuitive observation that remains to be resolved is that while ROS potentially drives the mitochondrial retrograde response, it also induces oxidative stress affecting polyploid yeast growth. ROS scavenging with reducing agents improved both mitochondria biogenesis (i.e., mt-DNA amount, Fig. 2d-f) and polyploid yeast survival under respiratory growth (Fig. 4l), suggesting that there is a fine balance between oxidative stress and physiological ROS signaling in response to polyploidy. We anticipate that future studies will identify the types of ROS species and their intracellular sources and decipher what ROS species are beneficial or inimical to polyploid yeast proliferation.

In conclusion, our study identifies sub-linear mitochondrial scaling with cell volume as a fundamental consequence of polyploidy. While cell size increases proportionally with genome content, mitochondrial biogenesis, including mitochondrial mass, network architecture, protein abundance, and mt-DNA content scales disproportionally. This alters mitochondria functions and induces a ROS-driven retrograde signaling mechanism to sustain metabolic homeostasis and cell proliferation under increased metabolic demands (Fig. 6i). These results establish mitochondrial scaling as a critical determinant of polyploid cell fitness and reveal retrograde signaling as an adaptive mechanism that buffers the metabolic consequences of genome duplication. Our findings shed light on how whole genome duplication may be linked to mitochondrial homeostasis and cellular fitness, which may be particularly relevant for metabolic regulation in cancer where whole genome duplication is a frequent phenomenon.

## Methods

### Yeast strains and growth conditions

All experiments were performed in the BY4741 genetic strain background. Isogenic polyploid yeast stains were constructed as previously described^11^. The list of strains used is provided in Supplementary Table 1. Gene deletion mutants were either obtained as gifts (publicly available from EUROSCARF) or created using a homologous recombination-based gene deletion approach involving amplification and transformation of antibiotic resistance markers into yeast cells using a standard lithium-acetate-polyethylene-glycol-based method^60, 95^. Created deletion strains were confirmed by polymerase chain reaction (PCR) on isolated genomic DNA using primers that anneal 100-200 bp upstream of the start codon and within the gene of interest. Where indicated, yeast strains were also transformed with constructs that integrate into the HO locus or plasmids harboring amino acid marker genes (Supplementary Table 2). Yeast cells were grown in yeast peptone (YP; 1% yeast extract, 2% bacto-peptone) or synthetic drop-out (S) media or agar plates supplemented with 2% glucose, galactose, glycerol, acetate, lactate or 3% ethanol. YPD supplemented with the appropriate antibiotic was used for selection of gene deletion mutants, while SC media lacking the appropriate amino acids was used for plasmid selection. Cells were incubated at 30°C with or without shaking at 140 rpm.

### Yeast growth assays

Growth assays were performed by drop dilution or growth curves. Overnight grown starter cultures were diluted to optical density (OD_600_) of 0.5 and cultured till OD_600_ of 0.8 – 1. One milliliter of 0.1 OD_600_ unit of cells resuspended in double-distilled water (ddH_2_O) was obtained to perform a dilution series of 1 in 10. Five microliters of cell suspension were spotted onto agars and incubated for at least 48 hours. For growth curves, 1 OD_600_ unit of cells were diluted to OD_600_ of 0.1 in 100 μl per well of a 96-well plate. The plate was sealed with a Breathe-Easy adhesive and incubated in a SPECTROstar Nano plate reader (BMG Labtech, Ortenberg, Germany) at 30°C with orbital shaking at 500 rpm. The OD_600_ of each well was recorded at every 10 min for 72 h. All experiments were performed in triplicate.

### Mating of yeast to generate triploid and tetraploid strains

Parental haploid and diploid yeast of Mat a and Mat α mating types^11^, as well as variants lacking *RTG2*, were mated by growth of both the Mat a and Mat α types together in YP media supplemented with 2% glucose. Alternatively, Mat a and Mat α cells were crossed on YPD plates^59^. Mating was verified by confocal microcopy via the visualization of shmoo formation. Mated cells harboring antibiotic resistance markers were selected by growth on YPD plates supplemented with the appropriate antibiotics. The resultant strains were confirmed via PCR with primers designed to amplify the gene or deletion cassette, and ploidy was ascertained via flow cytometry.

### Human cell lines and culture conditions

Post-tetraploid human cells were derived from the near-diploid human colon carcinoma cell line HCT116 (2N, ATCC No. CCL-247), with (i.e., p53^-/-^) or without (i.e., p53^+/+^) knockout of the p53 gene via CRISPR/Cas9, by inducing cytokinesis failure using 0.75 µM of the actomyosin inhibitor dihydrocytochalasin D (or DCD, Sigma) for 18 h. Cells were harvested, washed using phosphate buffered saline (PBS), sorted according to size by FACS and clonally expanded in drug-free medium by limiting dilution in a 96-well format. Additionally, a population of spontaneously arising tetraploid cells (i.e., Spon) were obtained from parental HCT116 cells via sorting by FACS^7^. Tetraploidy was validated by DNA content analysis using flow cytometry^5, 55^. All cells were cultured in Dulbeccós Modified Eagle Medium (DMEM; Gibco) supplemented with 10% Fetal Bovine Serum (FBS) and 1% Penicillin/Streptomycin and incubated at 37°C with 5% CO_2_ atmosphere.

### Ploidy confirmation of yeast and human cell lines by flow cytometry

To analyze DNA content, 1 OD_600_ unit of exponentially growing yeast cells was harvested and fixed with 70% ethanol at 4°C overnight with overhead rotation. Cells were then washed with ddH_2_O and treated with 0.1 mg/ml RNase A prepared in 0.5 ml 50 mM sodium citrate and incubated at 37°C for 2 h to digest RNA. Subsequently, cells were resuspended in 500 μl FxCycle™ PI/RNase staining solution (Life Technologies) and incubated in the dark at room temperature (RT) for 1 h. Cultured human cells were harvested, fixed and permeabilized at RT for 15 min. After washing in 1x PBS, the cell pellets were resuspended in PBS/DAPI solution supplemented with RNase A and incubated for 5 min at RT in the dark. All samples were analyzed on an Attune™ Flow Cytometer (Thermo Fisher Scientific) and data analysis was performed using the FlowJo™ software, version 10.

### Analysis of total proteome of tetraploid and parental human cells by tandem mass tag (TMT)-based quantitative proteomics

Cultured cell pellets (1 × 10^6^ cells) were snap-frozen and lysed in lysis buffer consisting of 1% SDS in 100 mM triethylammonium bicarbonate (TEAB). Lysates were sonicated in a water-bath sonicator using repeated cycles of 30 s sonication and 30 s cooling to reduce sample viscosity and shear genomic DNA. Following centrifugation at 8,000 × g for 15 min at 4°C, protein concentrations were determined using a BCA assay (Thermo Fisher Scientific). For each sample, 50 μg of total protein was adjusted to a final volume of 100 μl in 100 mM TEAB. Proteins were reduced with 10 mM tris(2-carboxyethyl)phosphine (TCEP) for 1 h at 55°C and alkylated with 18.75 mM iodoacetamide for 30 min at RT in the dark. Proteins were precipitated overnight at −20°C by addition of six volumes of ice-cold acetone. Protein pellets were collected by centrifugation, resuspended in 100 mM TEAB and digested overnight at 37°C with sequencing-grade trypsin at an enzyme-to-protein ratio of approximately 1:50.

For tandem mass tag (TMT)-based quantification, peptides were labeled using the TMT 6-plex Isobaric Label Reagent Set (Thermo Fisher Scientific). TMT reagents were dissolved in anhydrous acetonitrile according to the manufacturer’s instructions. Fifteen micrograms of peptides from each sample were incubated with the corresponding TMT label for 1 h at RT. Labeling reactions were quenched by addition of 5% hydroxylamine and incubated for 15 min. Equal amounts of labeled samples belonging to one biological replicate were subsequently pooled and dried by vacuum centrifugation.

### Peptide fractionation and mass spectrometry analysis

Prior to mass spectrometric analysis, pooled TMT-labeled peptide samples were fractionated using the Pierce High pH Reversed-Phase Peptide Fractionation Kit (Thermo Fisher Scientific) according to the manufacturer’s instructions. Briefly, dried peptide pellets were dissolved in 0.1% trifluoroacetic acid (TFA), loaded onto conditioned fractionation spin columns and separated into eight fractions using stepwise elution with increasing concentrations of acetonitrile in triethylamine-containing buffers. Fractions were collected individually and dried by vacuum centrifugation before LC–MS/MS analysis.

### Proteomics data processing

Protein group output files generated by MaxQuant were imported into R for downstream analysis. Reporter ion intensities were extracted from the corrected TMT reporter intensity columns. Protein groups annotated as potential contaminants, reverse database hits, or proteins identified only by site were removed prior to analysis. Protein identifiers were derived from the majority protein accession for each protein group. Sample annotations were matched to reporter intensity channels to generate expression matrices for downstream statistical analyses. Protein annotation, including UniProt accession numbers, Ensembl gene identifiers, approved gene symbols, chromosome assignments and gene names, was incorporated using BioMart-derived annotation tables.

Reporter intensities equal to zero were treated as missing values and converted to NA prior to normalization. Quantitative protein intensities were log2-transformed. To account for inter-replicate variability, batch effects associated with biological replicate were removed using the *removeBatchEffect* function implemented in the *limma* package. Subsequently, cyclic loess normalization was applied using the *limma* package to reduce systematic technical variation between samples. Data quality was assessed by boxplot visualization, principal component analysis (PCA), and hierarchical clustering represented as heatmaps. PCA was performed using centered and scaled normalized protein abundance values after removal of proteins containing missing values.

### Differential protein abundance analysis

Differential protein abundance was assessed independently for each experimental dataset using protein-wise linear regression models implemented in R. For each protein, normalized log2 abundance values were modeled as a function of treatment group using ordinary least-squares regression. Near diploid parental control samples served as the reference level, and treatment-specific regression coefficients were extracted to quantify abundance changes relative to controls. Comparisons were performed for all treatment conditions, including DCD induced und Spontaneous arising tetraploid samples. Resulting p-values were adjusted for multiple testing using the Benjamini–Hochberg procedure, and adjusted p-values were used to control the false discovery rate (FDR). Proteins were annotated with Ensembl gene identifiers and chromosome information using the *biomaRt* package.

### Analyses of additional proteomics data from budding yeast

Quantitative proteomics data for budding yeast strains with increasing ploidy was obtained from^11^ and data for isogenic aneuploid budding yeast was obtained from^46^. Published datasets from^12^ describing protein scaling behavior during G1 arrest and in cell-size mutants were used for comparative analyses. All computational analyses were performed in R (v4.3) and Python using packages for data processing, statistical analysis, and visualization, including tidyverse, ggplot2, limma, pheatmap, NumPy, SciPy, and matplotlib. Proteins were grouped into functional categories using gene ontology biological process (GOBP) and cellular component (GOCC) annotations. Classification was performed using regular-expression matching against GO annotation columns using grepl() with case-insensitive matching. Protein abundance tables were reshaped into log-format data frames using pivot_longer() for statistical visualization and group-wise comparisons. Ploidy states were encoded as ordered factors (1N, 2N, 3N, 4N). Expression values were converted to numeric values prior to downstream analysis. Differential protein abundance between ploidy states was evaluated using precomputed statistical outputs derived from MaxQuant analysis tables.

To quantify proteome scaling across increasing ploidy states, protein-specific scaling slopes were calculated using linear regression. For each protein, log2 fold changes relative to haploid cells were extracted: *FC*_2*N*_, *FC*_3*N*_, *FC*_4*N*_. The haploid state was assigned a baseline value of zero: *y* = [0, *FC*_2*N*_, *FC*_3*N*_, *FC*_4*N*_]. Protein scaling was then modeled using linear regression: *y* = *β*_0_ + *β*_1_*x*, where *x* = [1,2,3,4] represents ploidy level. Kernel density estimation plots were generated to visualize global proteome scaling distributions and mitochondrial protein scaling behavior across ploidy states. Protein abundance distributions were median scaled to experimentally observed fold change medians prior to plotting. Density curves were generated separately for total proteome abundance and mitochondrial protein abundance. Observed medians were overlaid as vertical reference lines. Density distributions and scaling curves were visualized using geom_density(), geom_point(), and nonlinear regression trend lines.

The regression coefficient (*β*_1_) was extracted and defined as the ploidy-dependent protein scaling slope. Positive slopes indicate increasing abundance with ploidy, whereas negative slopes indicate progressive dilution. To separate ploidy-dependent effects from cell-size-dependent effects, experimentally measured median cell volumes were incorporated into regression analyses. Median cell volumes for polyploid yeast cells were; 1N: 48.0 fL, 2N: 82.9 fL, 3N: 146.6 fL, 4N: 181.7 fL^11^. The median volumes of isogenic aneuploid yeast cells were obtained from^47^. Protein scaling relative to cell size was modeled using linear regression against measured cell volumes. The resulting regression coefficient was defined as the size-dependent protein scaling slope. Ploidy-dependent and size-dependent slopes were compared using scatter plot analyses and Pearson correlation.

To evaluate deviations from theoretical linear proteome scaling, power-law scaling models were fitted using nonlinear least squares. Median protein abundance values across ploidy states were compared against expected scaling behavior. Scaling relationships were modeled using a power-law equation: *y* = *a* × *x^b^*, where *x* corresponds to ploidy level and *y* to median protein abundance. The exponent reported by^12^ was used as a reference scaling model. Two-dimensional GO annotation enrichment analyses were performed by comparing annotation enrichment scores calculated from the polyploidy dataset^11^ against enrichment scores reported for G1-arrested cells and cell-size mutant datasets^12^.

Heatmaps were generated for functional protein groups whose gene sets were manually curated and filtered using dplyr. Expression matrices were generated from log2 fold change values across ploidy states. Replicate-specific fold changes were averaged across experiments. For each protein, log2 fold change values and associated q-values were extracted from contrasts comparing 2N vs 1N, 3N vs 1N, and 4N vs 1N. Heatmaps were visualized using the pheatmap package().

### Genomic DNA / RNA isolation, qPCR, and RT-qPCR

Exponentially growing yeast cultures were harvested and lysed in RNase-free tubes containing acid washed nuclease free glass beads using the FastPrep-24™ 5G disruptor (MP Biomedicals, Heidelberg, Germany) under the following conditions: 3 cycles of 20 s at speed 6 m/s and 2 min break in between each cycle. Genomic DNA (gDNA) and RNA were isolated using the Qiagen QIAamp DNA and RNeasy kits, respectively, according to the manufacturer’s instructions. Gene specific DNA levels were quantified from 10 ng of total gDNA using gene-specific primers. To quantify mRNA levels, cDNA was synthesized from 500 ng total RNA using iScript Advanced cDNA synthesis kit (Roche). Quantitative PCR was performed using gene specific primers and SsoAdvanced Universal SYBR Green Supermix (Bio-Rad) under the following thermocycling conditions: initial denaturation at 95°C for 3 min, followed by 40 cycles of 15 s denaturation at 95°C and amplification at 58°C for 30 s. A final heating gradient was performed from 58°C to 65°C for 5 s and then to 95°C at 0.5 °C/cycle. In all instances *ACT1* was used as a housekeeping gene. To analyze mitochondrial DNA (mt-DNA) copy number in human cells, total DNA was extracted from cultured cells using the QIAamp DNA Mini Kit according to the manufacturer’s instructions. Here, the nuclear encoded ß2 microglobulin gene was used to normalize the levels of mitochondria encoded genes and samples were run under the following conditions: initial denaturation for 5 min at 95°C followed by 45 cycles of 5 s at 95°C, 15 s at 55°C, 15 s at 72°C and 1 s at 78°C^63^. All qPCRs were run on the C1000 Touch Thermal Cycler (Bio-Rad). All primers used in this study are provided in Supplementary Table 3.

### Analysis of protein aggregation

Yeast strains transformed with an Hsp104-GFP construct were grown to mid-log phase in S-medium supplemented with 2% galactose at 30°C. In a control heat-shock reaction, cells were further incubated at 37°C for 2 h. Cells were harvested, washed with ddH_2_O, and visualized using the Leica DMi8 microscope with an external LED8 light source, a Leica DFC9000GTC camera and the HC PL APO 100x/1.44 oil objective. Z-stacked images were acquired using an HC PL APO100x/1.44 Oil UV objective with Immersion Oil Type A 518 F and analyzed with the LAS X software and ImageJ. A DFT51010 filter cube for GFP excitation at 475 nm at 540 MHz as quality mode at 16-bit and saved in the lif format. Brightfield images were acquired using the transmitted light detector. Images were processed via Fiji and the Leica Applications Suite X program (Ver 3.7.4).

### Analysis of mitochondrial network

Yeast strains transformed with a genome integrating Su9-mNeonGreen construct were grown to mid-log phase in S-medium supplemented with 2% galactose at 30°C. Images were taken with the Leica DMi8 microscope with an external LED8 light source, a Leica DFC9000GTC camera and the HC PL APO 100x/1.44 oil objective. Z-stacked images were acquired using an HC PL APO100x/1.44 Oil UV objective with Immersion Oil Type A 518 F and analyzed with the LAS X software and ImageJ. A DFT51010 filter cube for mNeonGreen excitation at 475 nm at 540 MHz as quality mode at 16-bit and saved in the lif format. Brightfield images were acquired using the transmitted light detector. Human cells were cultured on coverslips coated with poly-L-lysine and grown until about 80% confluency. Cells were washed with PBS, fixed using 2% paraformaldehyde (PFA)/ 2% sucrose solution, permeabilized with 0.1% Triton X-100 and blocked using 3% BSA in PBS supplemented with 0.1% Tween-20 (i.e., 3% BSA/PBST). Mitochondrial network was visualized by TOMM20 immunostaining and DAPI was used as a nuclear counterstain^63^. Images were acquired using a semi-automated Zeiss Axio Observer Z1 (Oberkochen, Germany) equipped with an ASI MS-2000 stage (Applied Scientific Instrumentation, Eugene, USA), a CSU-X1 spinning disk confocal head (Yokogawa), a Laser Stack with selectable lasers (Intelligent Imaging Innovations, Denver, USA) and the Cool-Snap HQ camera (Roper Scientific). The 63x oil objective was used under the control of the Slidebook6 software (Intelligent Imaging Innovations, Denver, CO). For budding yeast mitochondrial area was evaluated in ImageJ using macro based on thresholding, created in-house^50^. ROIs defining the cells were manually selected for each image and the area occupied my mitochondria calculated using the analyze particle function in ImageJ. For human cells, thresholding was performed on collapsed z-stack images using the “default” method, with the lower threshold manually adjusted to a fixed value between 40 and 60 (ensuring consistent segmentation across cell lines). Mitochondria were segmented using polygon ROIs drawn around individual cells guided by the DAPI signal. The percentage of cell surface area occupied by mitochondria was recorded for each ROI^56, 57^.

### Analysis of mitochondrial mass and membrane potential

Overnight grown starter cultures were diluted to OD_600_ of 0.1 and grown in appropriate media for 16-24 h. Subsequently, 0.5 OD_600_ units of cells were harvested, resuspended in 5 ml 1x PBS and aliquoted per ml into 1 ml Eppendorf tubes. Three tubes were processed as follows: one untreated, one treated with 2 µM tetramethylrhodamine ethyl ester perchlorate (TMRE; Invitorgen), and another treated with 250 nM nonyl acridine orange (NAO; Invitrogen) and incubated at 37 °C for 30 min. The samples were analyzed by flow cytometry on an Attune™ Flow Cytometer using appropriate filters. Data analysis was performed using the FlowJo™ software, version 10. The fluorescence intensity of the untreated sample was used for background correction. Mitochondrial mass was determined from mean NAO intensities and mitochondrial membrane potential (ΔΨ_m_) was calculated by normalizing the mean fluorescent intensity of TMRE with NAO intensity.

### Analysis of mitochondrial DNA nucleoids

Exponentially growing yeast cells in YP media supplemented with 2% glucose were treated with 2.5 µg/ml of DAPI and grown further for 1 h at RT in the dark. Cells were then washed with 1x PBS and visualized by microscopy using Zeiss Z1 Axio Observer microscope equipped with spinning disc head (Yokogawa, Hersching, Germany), CoolSNAP-HQ2 and CoolSNAP-EZ CCD cameras (Photometrics, Intelligent Imaging Innovations, Inc., Göttingen, Germany), Plan Apochromat 63x magnification oil objective, 40x magnification air objective or 20x magnification air objective, epifluorescence X-Cite 120 Series lamp and lasers: 473, 561 and 660 nm (LaserStack, Intelligent Imaging Innovations, Inc., Göttingen, Germany).

### Mitochondria isolation

Isolation of mitochondria was performed as previously described^96^. Briefly, 12 OD_600_ units of exponentially growing yeast cells in YP media supplemented with 2% galactose (2% glucose for *rho*^0^ cells) were harvested at 2000 g for 5 min at RT. Subsequently, cells were washed with 1.2 M sorbitol and treated with 2 ml/g wet weight of MP1 buffer (100 mM DTT, 10 mM Tris pH unadjusted) at 30°C for 10 min. Cells were washed again with 1.2 M sorbitol, resuspended in 6.7 ml/g wet weight of MP2 buffer (20 mM KPi buffer pH 7.4, 1.2 M sorbitol, 3 mg/g wet weight zymolyase 20T) and incubated at 30°C for 1 h. Spheroplasts were pelleted at 2000 g for 5 min at 4 °C and resuspended in 13.4 ml/g wet weight of ice-cold homogenization buffer (10 mM Tris pH 7.4, 1 mM EDTA pH 8, 0.2% fatty acids free bovine serum albumin, 1 mM PMSF, 0.6 M sorbitol). Spheroplasts were then homogenized by manual application of 10 strokes using a glass potter on ice. To collect mitochondria, cell debris was removed at 1500 g for 5 min, and supernatant further centrifuged at 12000 g for 12 min. Mitochondria were resuspended in 1 ml of ice-cold SH-buffer (0.6 M sorbitol, 20 mM HEPES pH 7.4) and aliquoted in equal volumes for subsequent use.

### Analysis of mitochondrial translation *in organello*

Mitochondrial *in organello* translation (ioTL) was performed as previously described ^96^. Briefly, equal aliquots of isolated mitochondria were incubated at 30°C and shaking at 600 rpm for 10 min in 1.5x ioTL buffer (20 mM HEPES/KOH pH 7.4, 15 mM KPi, 0.6 M Sorbitol, 150 mM KCl, 12.66 mM MgSO_4_, 12 μg/mL of all amino acids except methionine, 7.5 mM Phosphoenolpyruvate, 6 mM ATP, 0.75 mM GTP, 5 mM α-Ketoglutarate, 10 μg/mL Pyruvate kinase) supplemented with 1 μL of a 11 μCi solution of ^35^S-methionine. Radioactive labelling was quenched by the addition of 8 mM cold non-radioactive methionine and the reaction stopped via the addition of 1 mL ice-cold SH buffer. Samples were pelleted at 20000 g for 10 min at 4°C, resuspended in 25 µl of Laemmli buffer and analyzed by SDS-PAGE and autoradiography.

### Mitochondrial oxygen consumption assay

Rate of oxygen consumption of isolated mitochondria was measured using two different approaches. Firstly, mitochondria were isolated from 12 OD_600_ units of cells, and equal aliquots were loaded into an oxygen sensor microplate containing SH buffer at 25°C, in a 96-well format. Oxygen consumption was initiated by the addition of 2 mM NADH and measured using the FirePlate-O_2_ oximeter (PyroScience, Germany). Secondly, the amount of mitochondria was equally adjusted by measuring protein concentration using the Bradford assay. Oxygen consumption capacity was measured using 50 µg of intact mitochondria as previously described^97^. Briefly, mitochondria were incubated in SH buffer supplemented with 1 mM MgCl_2_ and loaded into the chamber of a Clark-type oxygen electrode connected to a computer-operated Oxygraph control unit (Hansatech Instruments, Germany) at 25°C with continuous stirring. The upper (100%) and lower (0%) limits of dissolved oxygen were calibrated using distilled water and sodium dithionite, respectively. Oxygen consumption was initiated by the addition of 7 mM NADH. Data was recorded at intervals of 1 s using the Oxygraph Plus software (Hansatech Instruments, Germany).

### ROS measurements

Yeast cells were grown in YP media supplemented with 2% glycerol for 24 h at 30°C. Next, 1 OD_600_ units of cells were harvested, washed with 1x PBS and incubated with 5 µM CellROX Deep Red reagent for 30 min at 30°C in the dark. The staining solution was removed, cells washed again with 1x PBS and visualized by microscopy using Zeiss Z1 Axio Observer microscope equipped with spinning disc head (Yokogawa, Hersching, Germany), CoolSNAP-HQ2 and CoolSNAP-EZ CCD cameras (Photometrics, Intelligent Imaging Innovations, Inc., Göttingen, Germany), Plan Apochromat 63x magnification oil objective, 40x magnification air objective or 20x magnification air objective, epifluorescence X-Cite 120 Series lamp and lasers: 473, 561 and 660 nm (LaserStack, Intelligent Imaging Innovations, Inc., Göttingen, Germany).

### Intracellular hydrogen peroxide (H_2_O_2_) measurement using roGFP2Tsa2ΔC_R_ biosensor

The roGFP2Tsa2ΔC_R_ biosensor-based experiments were performed as previously described^59, 60^. Briefly, haploid and tetraploid yeast transformed with the p416TEF-roGFP2Tsa2ΔC_R_ probe were grown in synthetic media supplemented with 2% glucose and lacking uracil for plasmid selection. Cells transformed with an empty p416TEF plasmid were used as fluorescence background control. Overnight precultures were diluted and grown at 30°C with orbital shaking at 140 rpm until OD_600_ ∼ 3. Cells were harvested and resuspended in 100 mM MES/Tris pH 6.0 buffer and transferred in 200 µl aliquots at 1.5 OD_600_ units/well in a 96-well plate format. For each strain, 20 mM diamide and 100 mM DTT treated samples were included to calibrate full biosensor oxidation and reduction, respectively. RoGFP2 fluorescence was measured at excitation maxima of 400 nm and 488 nm, and an emission maximum of 510 nm using a BMG Labtech CLARIOstar fluorescence plate reader. After 5 cycles of initial fluorescence measurement, the biosensor response to H_2_O_2_ treatment was followed. The degree of roGFP2 oxidation (OxD_roGFP2_) and area under the curve (AUC) were calculated as previously described^59–62^.

### Statistical analysis

Statical analyses were performed using GraphPad Prism 10. Unless otherwise stated, two-tailed unpaired Student’s *t*-test was used to compare the means of two groups. For multiple comparisons of non-normally distributed data, one-way ANOVA on ranks (Kruskal-Wallis H Test) was performed followed by Dunn’s post hoc test. For proteomic heatmaps, statistical significance was evaluated using two-sided t-tests against zero followed by Benjamini– Hochberg correction. Proteins were classified as significantly upregulated or downregulated using thresholds of |log2 fold change| ≥ 1 and q-value < 0.05. Significance levels are displayed using star annotations. Group-wise statistical comparisons between functional protein categories and background proteomes were performed using Wilcoxon rank-sum tests. Statistical annotations were visualized using stat_compare_means() and geom_signif(). Summary statistics were calculated using grouped means and standard deviations.

## Acknowledgements

We thank Martin van der Laan for critical feedback on the final draft of this manuscript. We are grateful to Yury Bykov for the kind gifts of *rtg2Δ*, *hog1Δ* and *tor1Δ* strains from the EUROSCARF library and to Bruce Morgan for the roGFP2-based biosensor plasmids. We thank Florian Eckert, Selime Ayvaz, Adelajda Korcari and Izabela Korcari for their help with the experiments. This project was supported by grants from the Landesforschungsinitiative Rheinland-Pfalz BioComp to P.S.A. and Z.S., and from the German Research Foundation (DFG) (GRK2737-STRESSistance) to Z.S. Work in the laboratory of J.M.H. is funded by grants from the German Research Foundation (DFG) (GRK2737-STRESSistance), European Research Council (ERC AdG MitoCyto 101052639) and the German-Israeli Foundation (Nexus grant no. I-1561-412.13/ 2023).

## Author contributions

P.S.A. and Z.S. conceived the project. A.R.H. generated the near-tetraploid human cell lines and, together with M.R., performed proteomics and bioinformatic analyses. P.S.A., A.M.S., G.Y., A.H., and A.K. performed biochemical and cell biology experiments. P.S.A. performed data analyses. P.S.A., Z.S. and J.M.H. supervised the project. P.S.A. and Z.S. wrote the manuscript, and all authors critically reviewed and commented on the manuscript.

