## Supplementary for "Whole genome duplication uncouples cell volume scaling from mitochondrial homeostasis"

**This file contains:**

Supplementary Figures 1-6

Supplementary Tables 1-3

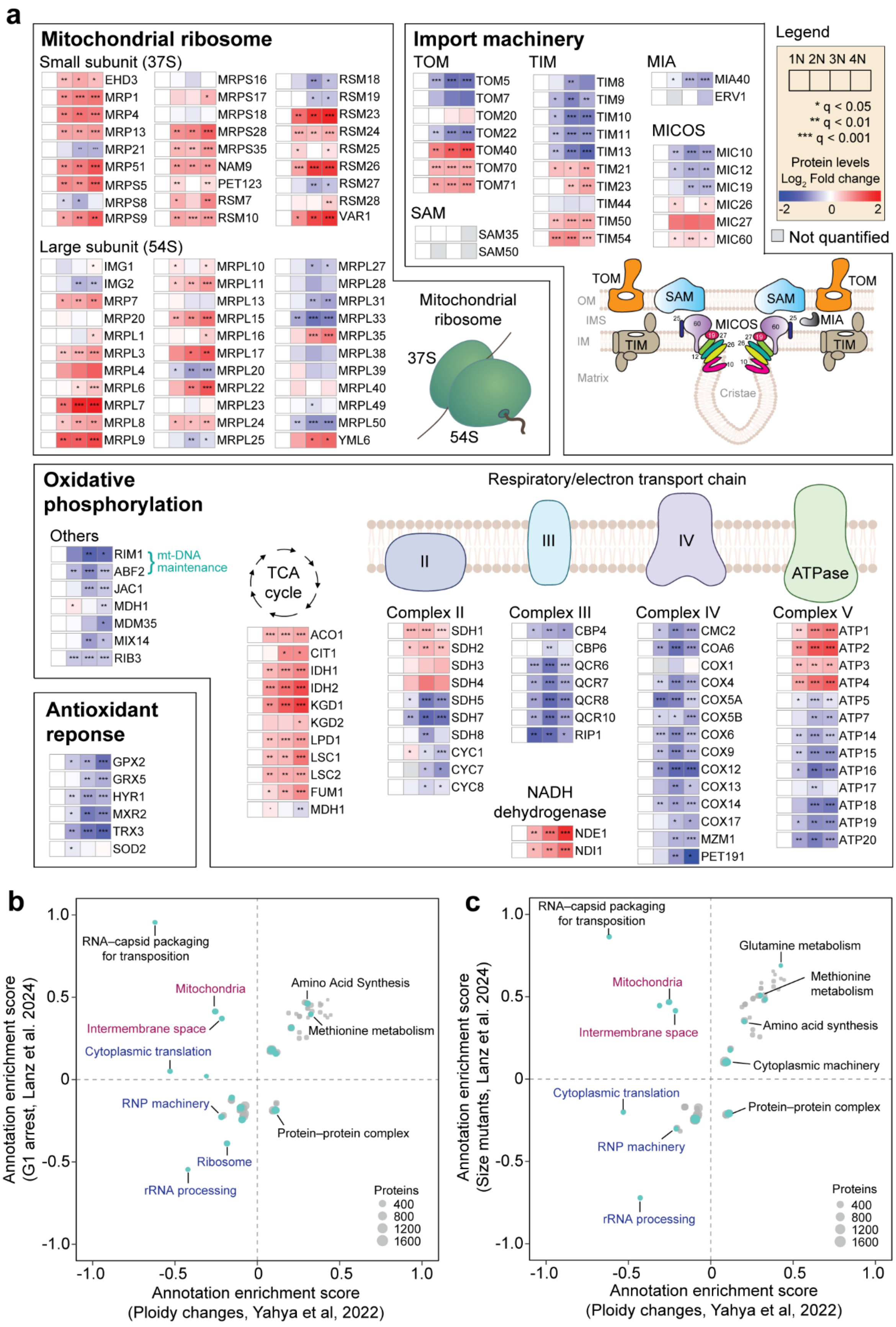
a
Mitochondrial ribosome
Small subunit (37S)
EHD3
MRP1
MRP4
MRP13
MRP21
MRP51
MRPS5
MRPS8
MRPS9
MRPS16
MRPS17
MRPS18
MRPS28
MRPS35
NAM9
PET123
RSM7
RSM10
RSM18
RSM19
RSM23
RSM24
RSM25
RSM26
RSM27
RSM28
VAR1
Large subunit (54S)
IMG1
IMG2
MRP7
MRP20
MRPL1
MRPL3
MRPL4
MRPL6
MRPL7
MRPL8
MRPL9
MRPL10
MRPL11
MRPL13
MRPL15
MRPL16
MRPL17
MRPL20
MRPL22
MRPL23
MRPL24
MRPL25
MRPL27
MRPL28
MRPL31
MRPL33
MRPL35
MRPL38
MRPL39
MRPL40
MRPL49
MRPL50
YML6
Import machinery
TOM
TOM5
TOM7
TOM20
TOM22
TOM40
TOM70
TOM71
SAM
SAM35
SAM50
TIM
TIM8
TIM9
TIM10
TIM11
TIM13
TIM21
TIM23
TIM44
TIM50
TIM54
MIA
MIA40
ERV1
MICOS
MIC10
MIC12
MIC19
MIC26
MIC27
MIC60
Legend
1N 2N 3N 4N
* q < 0.05
** q < 0.01
*** q < 0.001
Protein levels
Log2 Fold change
-2
0
2
Not quantified
Mitochondrial ribosome
37S
54S
OM
IMS
IM
Matrix
Cristae
Oxidative phosphorylation
Respiratory/electron transport chain
Others
RIM1
ABF2
mt-DNA maintenance
JAC1
MDH1
MDM35
MIX14
RIB3
TCA cycle
ACO1
CIT1
IDH1
IDH2
KGD1
KGD2
LPD1
LSC1
LSC2
FUM1
MDH1
II
III
IV
ATPase
Complex II
SDH1
SDH2
SDH3
SDH4
SDH5
SDH7
SDH8
CYC1
CYC7
CYC8
Complex III
CBP4
CBP6
QCR6
QCR7
QCR8
QCR10
RIP1
NADH dehydrogenase
NDE1
NDI1
Complex IV
CMC2
COA6
COX1
COX4
COX5A
COX5B
COX6
COX9
COX12
COX13
COX14
COX17
MZM1
PET191
Complex V
ATP1
ATP2
ATP3
ATP4
ATP5
ATP7
ATP14
ATP15
ATP16
ATP17
ATP18
ATP19
ATP20
Antioxidant reponse
GPX2
GRX5
HYR1
MXR2
TRX3
SOD2
b
RNA–capsid packaging for transposition
Mitochondria
Intermembrane space
Cytoplasmic translation
Amino Acid Synthesis
Methionine metabolism
RNP machinery
Protein–protein complex
Ribosome
rRNA processing
Proteins
400
800
1200
1600
Annotation enrichment score (G1 arrest, Lanz et al. 2024)
Annotation enrichment score (Ploidy changes, Yahya et al, 2022)
c
RNA–capsid packaging for transposition
Glutamine metabolism
Mitochondria
Methionine metabolism
Intermembrane space
Amino acid synthesis
Cytoplasmic machinery
Cytoplasmic translation
Protein–protein complex
RNP machinery
rRNA processing
Proteins
400
800
1200
1600
Annotation enrichment score (Size mutants, Lanz et al. 2024)
Annotation enrichment score (Ploidy changes, Yahya et al, 2022)

**Supplementary Figure 1: Proteomics analyses delineate cell size and ploidy-dependent changes in cellular pathways. a,** Heatmaps showing $log_2$ fold changes in expression of selected mitochondrial proteins relative to the haploid (1N) strain. Grey boxes indicate values not statistically quantified. * $q < 0.05$, ** $q < 0.01$, *** $q < 0.001$. Comparative 2D-annotation enrichment analyses of cellular pathways altered by **b,** increased cell size induced via G1 arrest[12] versus polyploidy-induced changes, and **c,** altered cell sizes based on gene deletions[12] versus polyploidy-induced changes, using the calculated protein slopes. Each dot is a significantly enriched annotation group, the dot size is determined by the number of proteins in each group, and the position of the dot is determined by the mean slope value of the protein group.

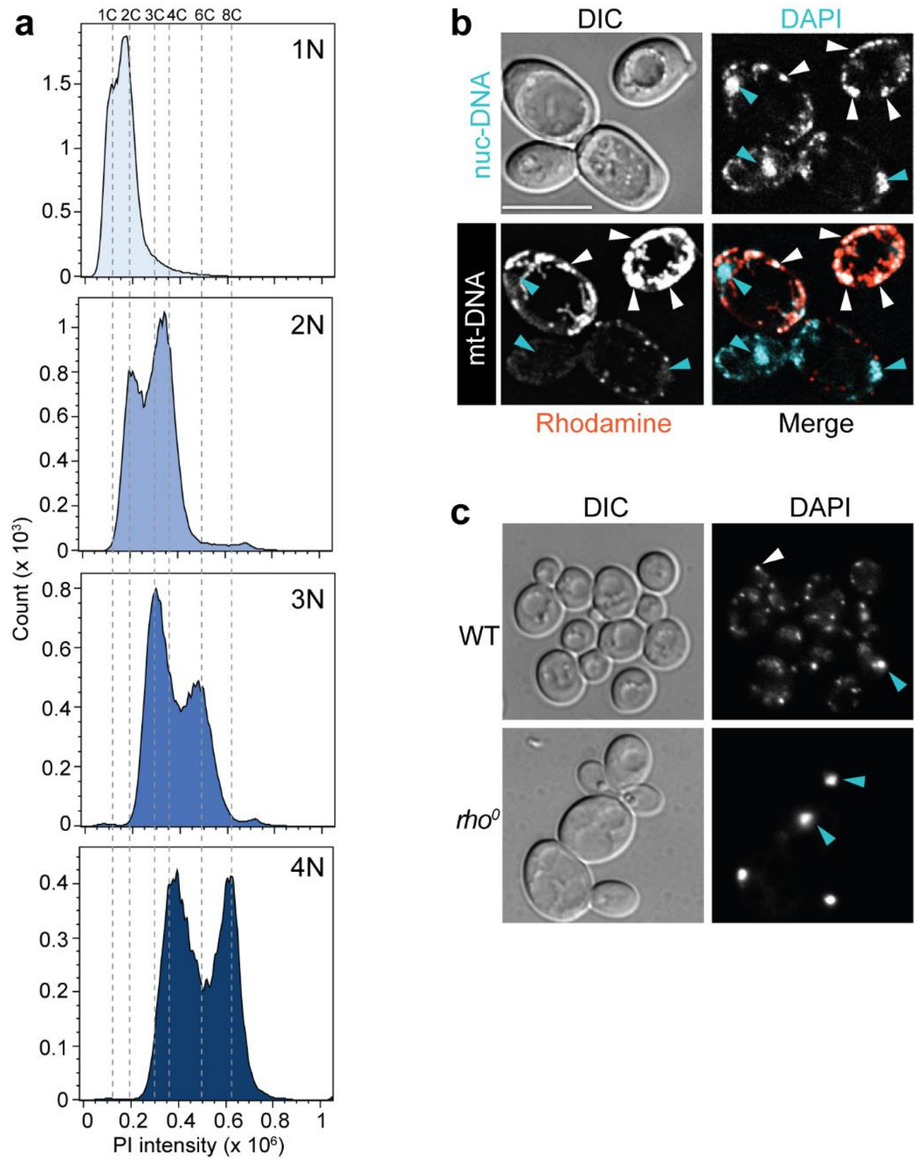


**Supplementary Figure 2: Confirmation of yeast ploidy and DNA staining. a,** Flow cytometry analyses of ploidy series of BY4741 yeast strain. **b,** Representative confocal images showing differential staining of nuclear (nuc-DNA) and mitochondrial (mt-DNA) DNA using DAPI. Rhodamine dye was used as a mitochondrial marker to localize mt-DNA foci. **c,** Representative confocal images showing exclusive nuclear DNA staining in haploid *rho*$^{0}$ cells lacking mt-DNA, compared to the WT showing both mt-DNA and nuclear DNA staining, and thereby clearly distinguishing mt-DNA from nuc-DNA foci. Scale bar is 5 µm. White arrowhead = mt-DNA, cyan arrowhead = nuc-DNA.

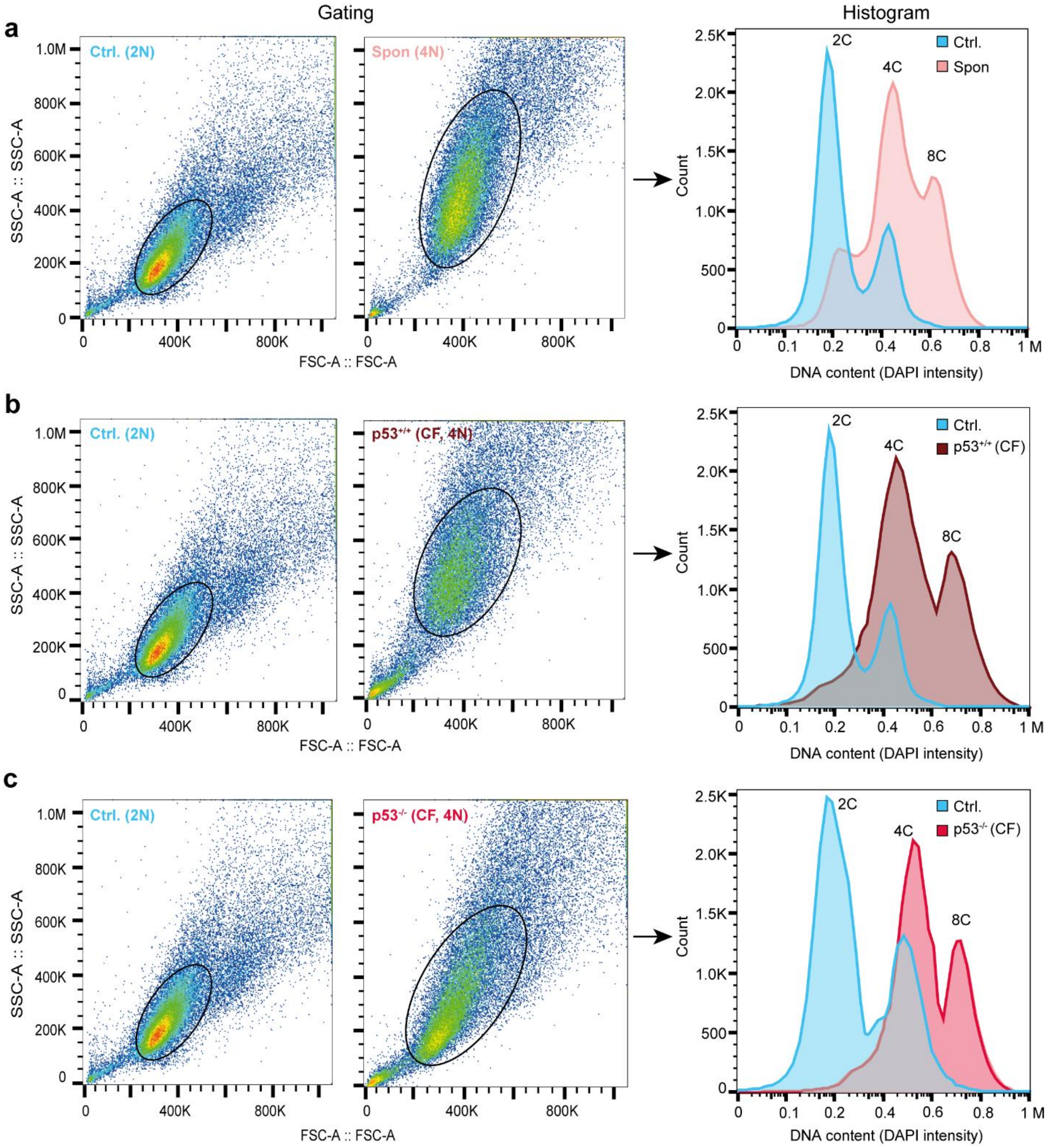


**Supplementary Figure 3: Confirmation of ploidy of HCT116 human cell lines.** Flow cytometry analysis confirms the ploidy of **a,** Spon, **b,** $p53^{+/+}$-CF and **c,** $p53^{-/-}$-CF cells, in comparison to the parental diploid (2N). The gating strategy and histogram of DNA content are shown.

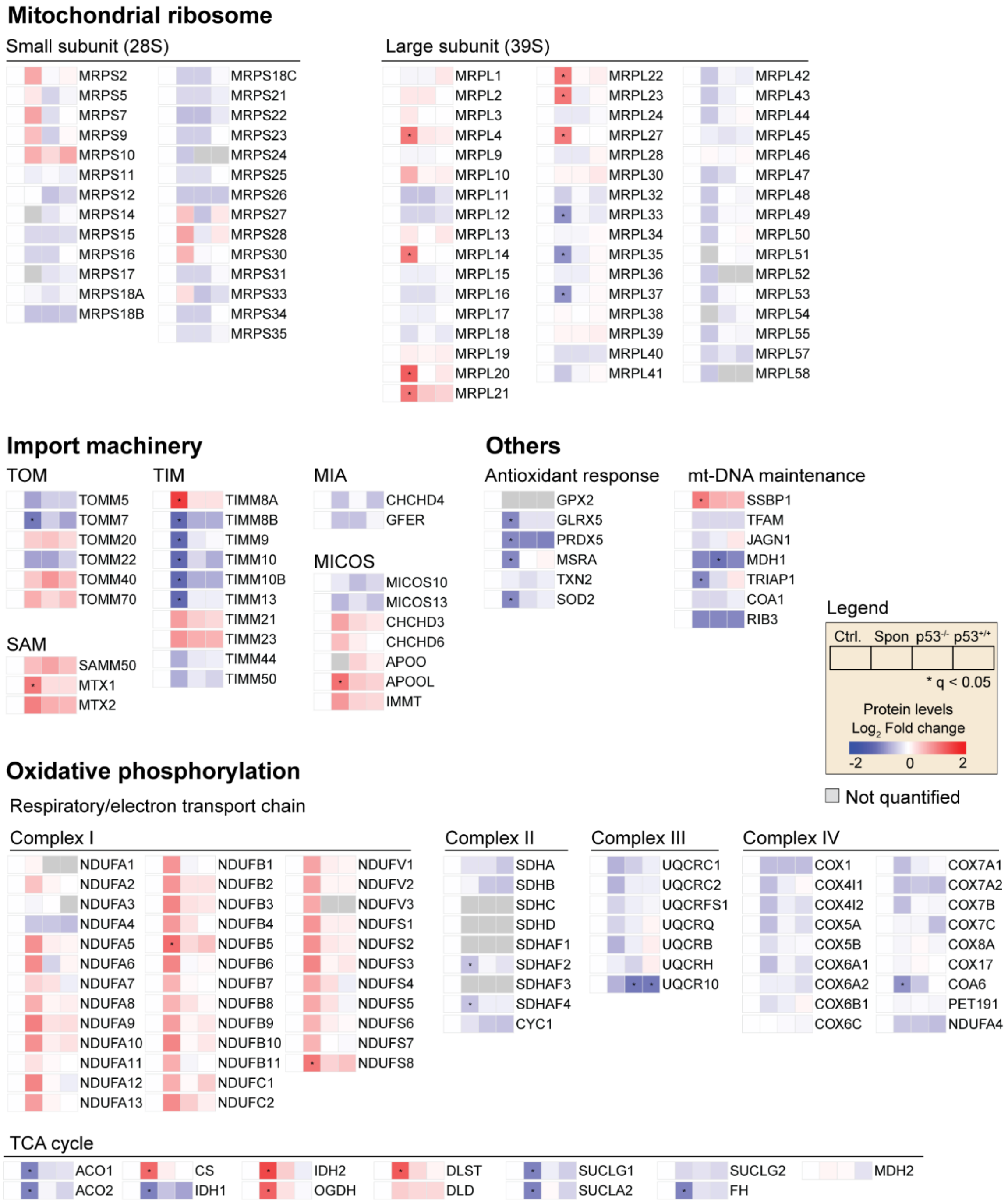


**Supplementary Figure 4: Proteomics analysis reveals changes in expression of mitochondrial proteins in response to polyploidy in human cells.** Heatmaps showing $\log_2$ fold changes in expression of selected mitochondrial proteins relative to the parental diploid. Grey boxes indicate values not statistically quantified. Components of complex V were not quantified from the proteomics analyses. * $q < 0.05$.

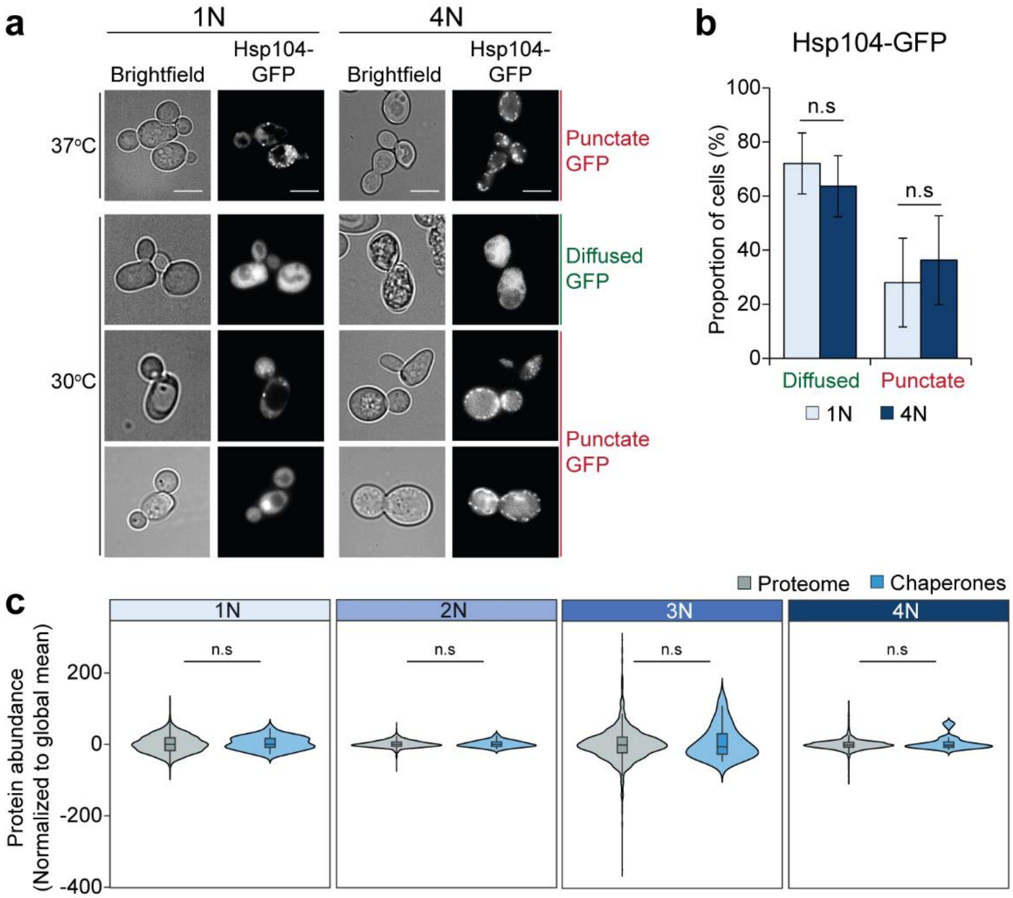


**Supplementary Figure 5: Polyploid budding yeast do not exhibit significant protein aggregation stress. a,** Representative confocal images showing Hsp104-GFP distribution in haploid (1N) and tetraploid (4N) yeast strains. Scale bar is 5 µm. Cells showing both diffused and punctate expression of Hsp104-GFP were evaluated. Heat shock induced at 37$^{o}$C was included as positive control for Hsp104-GFP puncta formation. **b,** Quantification of the proportion of cells with diffused or punctate Hsp104-GFP from (**a**). Data is shown as mean ± s.e.m. for n = 496 (1N) and n 269 (4N) cells from three independent experiments. *P*-values represent two-tailed unpaired Student's *t*-test. n.s = not significant. **c,** Comparative changes in abundance of chaperones to the global proteome in the yeast ploidy series. *P*-values show statistical comparison to the global proteome and are derived from two-sided Wilcoxon's ranks sum tests. n.s = not significant.

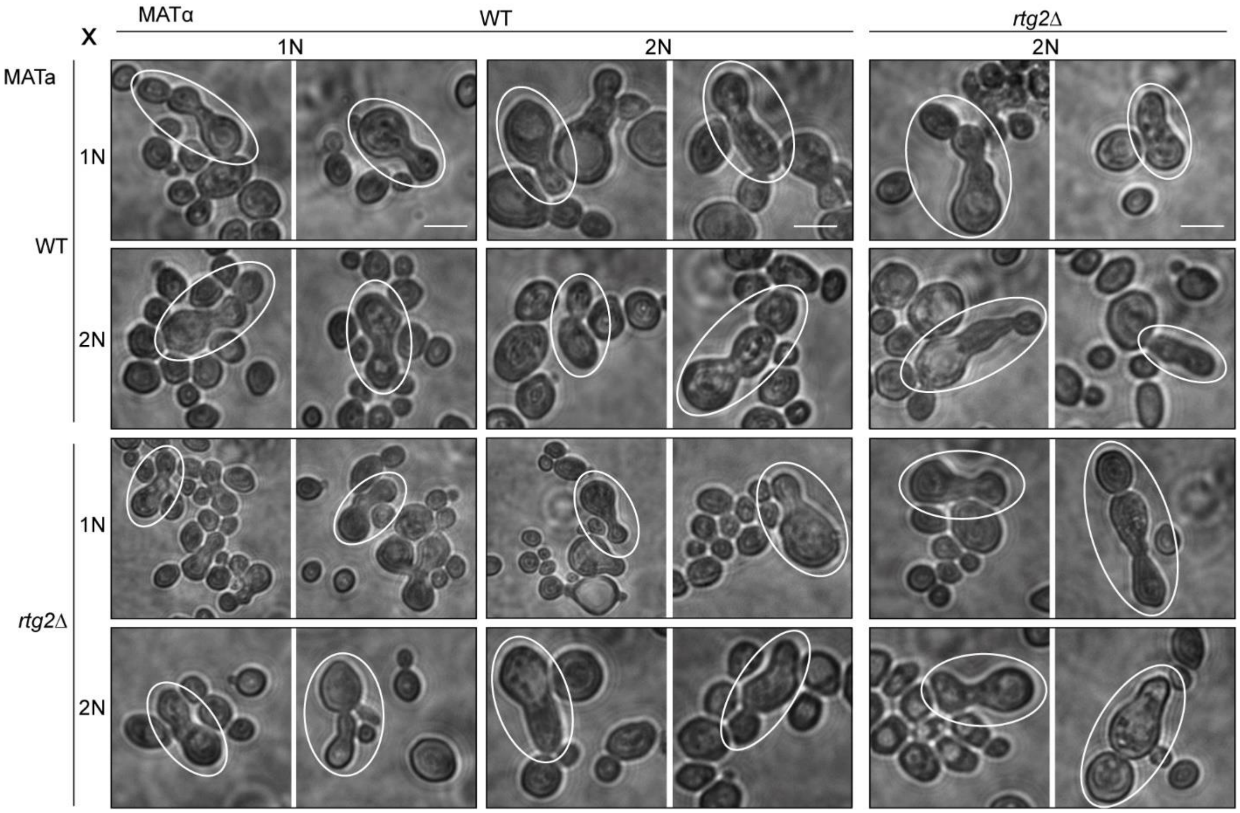


**Supplementary Figure 6: Impact of loss of *RTG2* on yeast mating and zygote formation.** Representative confocal images showing mating of haploid (1N) and diploid (2N) budding yeast either possessing (i.e., WT) or lacking *RTG2* (i.e., *rtg2Δ*). Images are shown in duplicates for each condition. Examples of zygotes formed from yeast mating are indicated with white circles. Scale bar is 5 µm.

**Table 1: Yeast strains used in this study**

| Name | Ploidy | Genotype |
|---|---|---|
| YZ944 | 1N | *MATa leu2Δ his3Δ ura3Δ LYS2* |
| YZ945 | 1N | *MATα leu2Δ his3Δ ura3Δ LYS2* |
| YZ946 | 2N | *MATa/a ura3Δ/ura3Δ leu2Δ/leu2Δ his3Δ/his3Δ LYS2/LYS2 met15Δ /MET15* |
| YZ947 | 2N | *MATα/α ura3Δ/ura3Δ leu2Δ/leu2Δ his3Δ/his3Δ LYS2/LYS2 met15Δ /MET15* |
| YZ949 | 3N | *MATa/a/a ura3Δ/ura3Δ/ura3Δ leu2Δ/leu2Δ/leu2Δ his3Δ/his3Δ/his3Δ LYS2/LYS2/LYS met15Δ/MET15/MET15* |
| YZ950 | 4N | *MATa/a/a/a ura3Δ/ura3Δ/ura3Δ/ura3Δ leu2Δ/leu2Δ/leu2Δ /leu2Δ his3Δ/his3Δ/his3Δ/his3Δ LYS2/LYS2/LYS2/LYS2 met15Δ/MET15/met15Δ/MET15* |
| YZ505 | 4N | *MATa/α/a/α leu2Δ0/leu2Δ0/leu2Δ0/leu2Δ0 ura3Δ0/ura3Δ0/ura3Δ0/ura3Δ0 his3Δ1/his3Δ1/his3Δ1/his3Δ1 Pds1-myc-LEU2* |
| YZ874 | 4N | *MATa/a/a/a ura3Δ0/ura3Δ0/ura3Δ0/ura3Δ0 leu2Δ0/leu2Δ0/leu2Δ0/leu2Δ0 his3Δ1/his3Δ1/his3Δ1/his3Δ1 met15Δ0/met15Δ0/MET15/MET1* |
| YZ1847 | 1N | *MATα Rho0* |
| YZ1931 | 1N | *MATa his3Δ1 leu2Δ0 met15Δ0 ura3Δ0 RTG2::kanMX4* |
| YZ1941 | 2N | *MATa/a leu2Δ/leu2Δ his3Δ/his3Δ ura3Δ/ura3Δ LYS2/LYS2 MET15/met15 RTG2::natNT2 RTG2::hphNTI* |
| YZ1942 | 2N | *MATα/α ura3Δ/ura3Δ leu2Δ/leu2Δ his3Δ/his3Δ LYS2/LYS2 met15Δ /MET15 RTG2::natNT2 RTG2::hphNTI* |
| YZ1943 | 3N | *MATa/α/α ura3Δ/ura3Δ leu2Δ/leu2Δ his3Δ/his3Δ LYS2/LYS2 met15Δ /MET15 RTG2::kanMX4 RTG2::natNT2 RTG2::hphNTI* |
| YZ1932 | 1N | *MATa his3Δ1 leu2Δ0 met15Δ0 ura3Δ0 TOR1::kanMX4* |
| YZ1947 | 1N | *MATa his3Δ1 leu2Δ0 met15Δ0 ura3Δ0 HOG1::kanMX4* |
| YZ1940 | 1N | *MATa his3Δ1 leu2Δ0 met15Δ0 ura3Δ0 TOR1::kanMX4 RTG2::natNT2* |
| YZ1948 | 1N | *MATa his3Δ1 leu2Δ0 met15Δ0 ura3Δ0 HOG1::kanMX4 RTG2::hphNTI* |
| YZ1949 | 1N | *MATa his3Δ1 leu2Δ0 met15Δ0 ura3Δ0 HOG1::kanMX4 TOR1::hphNTI* |

**Table 2: List of plasmids used in this study**

| Name | Comment |
|---|---|
| pRS303 Su9-ymNeonGreen | Genomically integrating plasmid. Linearized with PstI prior to transformation |
| pRS306 Su9-ymNeonGreen | Genomically integrating plasmid. Linearized with PstI prior to transformation |
| pYX142 Hsp104-GFP | Plasmid for analysis of protein aggregates |
| p416 TEF empty | Background control for cytosolic $H_2O_2$ measurement. Plasmid contains URA3 maker for selection after transformation |
| p416 TEF roGFP2-Tsa2Δ$C_R$ | Genetically encoded biosensor for cytosolic $H_2O_2$ measurement. Plasmid contains URA3 maker for selection after transformation |

**Table 3: Sequences of primers used in this study**

| Gene name | Forward sequence | Reverse sequence |
|---|---|---|
| *S.c COX3* | TTGAAGCTGTACAACCTACC | GCGCTCTAGCTCTGTGAGAT |
| *S.c RTG2* | GCCTCTTATCCAAAGGAGTT | TCCGTGCGAGCTTATTACCC |
| *S.c CIT2* | CGCCGACATTCAAAAGGACC | CTTGCGCTTGAGTTGGAACC |
| *S.c DLD3* | GTCAAGTGGGCGGTGTAGTT | ATCTCACCGTTGGGTAGCAC |
| *S.c ACT1* | CACCCTGTTCTTTTGACTGA | CGTAGAAGGCTGGAACGTTG |
| *H.s mt-D LOOP* | CTTCTGGCCACAGCACTTAAAC | GCTGGTGTTAGGGTTCTTTGTTTT |
| *H.s mt-ND1* | CCACCTCTAGCCTAGCCGTTTA | GGGTCATGATGGCAGGAGTAAT |
| *H.s mt-ATP6* | TAGCCATACACAACACTAAAGGACGA | GGGCATTTTTAATCTTAGAGCGAAA |
| *H.s* $ß_2M$ | GCTGGGTAGCTCTAAACAATGTATTCA | CCATGTACTAACAAATGTCTAAAATGGT |
